# The AKT2/SIRT5/TFEB pathway as a potential therapeutic target in atrophic AMD

**DOI:** 10.1101/2023.08.08.552343

**Authors:** Sayan Ghosh, Ruchi Sharma, Sridhar Bammidi, Victoria Koontz, Mihir Nemani, Meysam Yazdankhah, Katarzyna M. Kedziora, Callen T. Wallace, Cheng Yu-Wei, Jonathan Franks, Devika Bose, Dhivyaa Rajasundaram, Stacey Hose, José-Alain Sahel, Rosa Puertollano, Toren Finkel, J. Samuel Zigler, Yuri Sergeev, Simon C. Watkins, Eric S. Goetzman, Miguel Flores-Bellver, Kai Kaarniranta, Akrit Sodhi, Kapil Bharti, James T. Handa, Debasish Sinha

**Affiliations:** Department of Ophthalmology, University of Pittsburgh School of Medicine, Pittsburgh, PA, USA; Ocular and Stem Cell Translational Research Section, National Eye Institute, National Institutes of Health, Bethesda, MD, USA; Department of Cell Biology, Center for Biologic Imaging, University of Pittsburgh School of Medicine, Pittsburgh, PA, USA; Aging Institute, University of Pittsburgh School of Medicine, Pittsburgh, PA, USA; Department of Pediatrics, University of Pittsburgh School of Medicine, PA, USA; Wilmer Eye Institute, The Johns Hopkins University School of Medicine, Baltimore, MD, USA; Institut De La Vision, INSERM, CNRS, Sorbonne Université, Paris, France; Cell Biology and Physiology Center, National Heart, Lung, and Blood Institute, National Institutes of Health, MD, USA; Protein Biochemistry & Molecular Modeling Group, National Eye Institute, National Institutes of Health, Bethesda, MD, USA; Department of Ophthalmology, University of Colorado Anschutz Medical Campus, Aurora, Colorado, USA; Department of Ophthalmology, University of Eastern Finland and Kuopio University Hospital, Kuopio, Finland; Department of Molecular Genetics, University of Lodz, Lodz, Poland

## Abstract

Age-related macular degeneration (AMD), the leading cause of geriatric blindness, is a multi-factorial disease with retinal-pigmented epithelial (RPE) cell dysfunction as a central pathogenic driver. With RPE degeneration, lysosomal function is a core process that is disrupted. Transcription factors EB/E3 (TFEB/E3) tightly control lysosomal function; their disruption can cause aging disorders, such as AMD. Here, we show that induced pluripotent stem cells (iPSC)-derived RPE cells with the complement factor H variant [*CFH* (Y402H)] have increased AKT2, which impairs TFEB/TFE3 nuclear translocation and lysosomal function. Increased AKT2 can inhibit PGC1α, which downregulates SIRT5, an AKT2 binding partner. SIRT5 and AKT2 co-regulate each other, thereby modulating TFEB-dependent lysosomal function in the RPE. Failure of the AKT2/SIRT5/TFEB pathway in the RPE induced abnormalities in the autophagy-lysosome cellular axis by upregulating secretory autophagy, thereby releasing a plethora of factors that likely contribute to drusen formation, a hallmark of AMD. Finally, overexpressing AKT2 in RPE cells in mice led to an AMD-like phenotype. Thus, targeting the AKT2/SIRT5/TFEB pathway could be a potential therapy for atrophic AMD.

## Main text

Atrophic AMD is a leading cause of blindness among the elderly population worldwide^1^. Available treatments slow, but do not prevent, AMD progression. A key early pathogenic event is the degeneration of the RPE^2^. The RPE provides multiple sight-saving functions that depend on a high metabolic rate that also produces substantial and potentially toxic waste^3^. To maintain its health and photoreceptor viability needed for vision, the RPE relies heavily on lysosomal-mediated waste clearance and intact mitochondrial function^1, 3, 4^. Disruption of these processes leads to RPE dysfunction and contributes mechanistically to AMD development^1, 3, 4^.

Lysosomal biogenesis and function are under the control of the master regulator, transcription factor EB and/or E3 (TFEB and TFE3)^5^. TFEB and TFE3 belong to the MiTF/TFE (basic helix-loop-helix) transcription factor subfamily that regulates the transcription of genes in the lysosome-autophagy pathway and is, in turn, regulated by the mechanistic target of rapamycin, complex 1 (mTORC1)^5^. TFEB/TFE3 can also be regulated by mTOR-independent pathways such as AKT1/2 and calcium signaling^6–8^. In fact, Akt has been shown to phosphorylate TFEB at Ser467 and to repress TFEB nuclear translocation independently of mTORC1^6^. AKT2 is both an upstream regulator of mTORC1 and a downstream mediator for mTORC2^9,10^, and thus, plays a central role in regulating lysosomal function. Intriguingly, lysosomal function can be compensated by TFE3 activation in cells with diminished TFEB nuclear activity^11^. We found that upregulating AKT2 in TFEB null (*Tfeb* KO) mouse embryonic fibroblasts (MEFs) increased TFE3 phosphorylation (S321) and reduced expression of Coordinated Lysosomal Expression and Regulation (CLEAR) network genes, including Cathepsin D and L (Supplementary Fig. 1a, b). In addition, AKT2 overexpression in *Tfeb* KO cells induced TFE3 binding to 14-3-3 proteins, which is known to sequester TFE3 in the cytoplasm (Supplementary Fig. 1c). These results indicate that TFE3 can compensate for TFEB loss and mitigate lysosomal dysfunction.

AKT2 was previously reported to be upregulated in dysmorphic macular RPE cells from AMD donor globes^12, 13^. Moreover, in a well-characterized mouse model of lysosomal impairment that develops an AMD-like phenotype (the *Cryba1* cKO), AKT2 is increased in the RPE, which impairs lysosomal biogenesis^14, 15^. However, it is unknown whether dysfunctional AKT2 signaling directly contributes to AMD progression. To explore this question, we generated an RPE-specific *Akt2* knockin (KI) mouse, as previously described^16^, and found that CLEAR network genes were downregulated in the RPE while such changes were not observed in *Akt2* conditional knockout (cKO) RPE (Fig. 1a, b). *Akt2* KI mice developed an AMD-like phenotype, as indicated by an age-dependent accumulation of multiple autofluorescent foci compared to age-matched controls, as seen on fundus images (Supplementary Fig. 2a). By 10 months of age, the RPE in *Akt2* KI mice was morphologically heterogeneous with dysmorphic cells intermixed among the normal cobblestone shaped cells both in peripheral and central retina, while WT controls had uniform RPE morphology (Fig. 1c and Supplementary Fig. 2b). By 12 months, the dysmorphic RPE in *Akt2* KI retina had reduced apical microvilli length relative to age-matched WT mice (Fig. 1d). Moreover, as the mice age, further abnormalities in microvilli (disintegration) and photoreceptor outer segments (circular shape) were observed by transmission electron microscopy (TEM) images in 15 month old *Akt2* KI RPE; these were not seen in young mice (Supplementary Fig. 2c). Ezrin^17^, an important apically located linker protein critical for maintaining RPE differentiation and polarity, was significantly decreased in *Akt2* KI mice (Supplementary Fig. 2d). We also found decreased Ezrin-Radixin-Moesin binding Phosphoprotein 50 (EBP50) expression in 12-month-old *Akt2* KI retina (Fig. 1d). The RPE also accumulated lipid deposits that were confirmed by Perilipin-2 (PLIN2) staining^14^ of sections from 15-month-old *Akt2* KI mouse eyes (Supplementary Fig. 2e). The Bruch’s membrane of *Akt2* KI mice developed basal laminar deposits (BLamD) by 15 months of age, which were not seen in age-matched controls (Fig. 1e). Immunofluorescence studies of rhodopsin showed diffused staining and significant reduction in the expression of the protein in 12-month-old *Akt2* KI retina (Fig. 1d). These histologic, and corresponding molecular changes, correlated with decreased a- and b-wave amplitudes on electroretinograms (ERG) of old *Akt2* KI mice while no ERG changes were observed in age-matched WT control mice (Supplementary Fig. 2f, g). Collectively, these morphologic, ultrastructural, molecular, and physiologic studies suggest that activated AKT2 signaling directly contributes to an early atrophic AMD phenotype.

**Figure 1:**
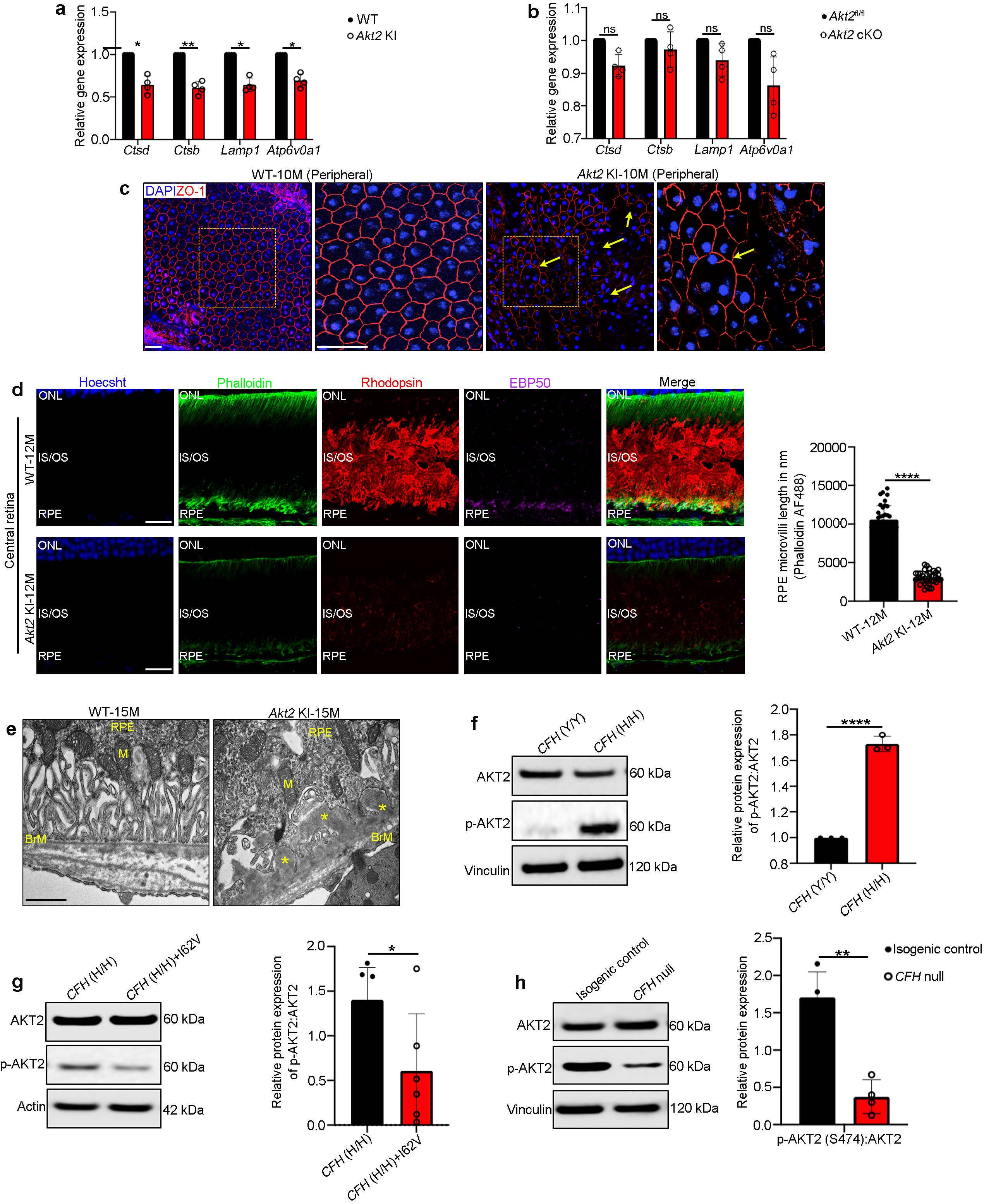
Akt2 upregulation triggers lysosomal dysfunction and an AMD-like phenotype in mice. Decline in expression of CLEAR network genes *Ctsd*, *Ctsb*, *Lamp1* and *Atp6v0a1* in (**a**) *Akt2* KI, but not in (**b**) *Akt2* cKO RPE cells. n=4. (**c**) Immunostaining with ZO-1 on RPE flatmounts from 10-month-old *Akt2* KI mice shows dysmorphic RPE cells in the peripheral regions (arrows in **c**), not seen in age-matched WT RPE. n=4. Scale bar= 50 μm (Zoomed Inset= 80 μm). (**d**) Immunofluorescence studies revealed noticeable decrease in EBP50 (magenta) and rhodopsin (red) staining in the RPE, as well as decreased microvilli length (green and graph) in retinal sections from12-month-old *Akt2* KI mice, compared to WT. n=4. (**e**) Transmission electron micrographs showing accumulation of basal laminar deposits in the RPE cells (asterisks in **e**) above Bruch’s membrane (BrM) in 15-month-old *Akt2* KI mice, but not in age matched WT RPE cells. M=mitochondria. n=5. Scale bar 600 nm. Western blot analysis showing (**f**) elevated p-Akt2:Akt2 ratio in iPSC-derived RPE cells from *CFH* Y402H risk allele [homozygous; *CFH* (H/H)] containing donors (with no AMD), relative to controls [*CFH* (Y/Y)] (n=3), and (**g**) reduction of the p-Akt2:Akt2 ratio when *CFH* I62V mutation is also present in CFH (H/H) cells (n=6) or (**h**) by complete knockout of *CFH* (*CFH* null) in iPSC-derived RPE cells. n=3. All values are Mean ± S.D. *P<0.05, **P<0.01, ****P<0.0001.

In addition to age and environmental factors such as smoking and sun exposure, a number of single nucleotide polymorphisms (SNPs) have been associated with elevated AMD risk^1^. Genome-Wide Association Studies (GWAS) have identified over 50 SNPs that confer AMD risk, many of which affect genes involved in the complement cascade^18, 19^. Among the complement genes implicated in AMD risk is the complement factor H (CFH)-CFH Receptor 5 (CFHR5) region on chromosome 1q32. CFH-CFHR5 is a major genetic risk locus for AMD, increasing the odds of disease by more than 2.5 fold among heterozygotes and 7.5 fold among homozygotes^18, 19^. Moreover, iPSC-derived RPE cells from donors with the CFH 402H risk allele have lysosomal functional defects^20^. We, therefore, set out to determine whether CFH risk alleles influence the regulation of lysosomal function by AKT2. To this end, we examined AKT2 levels in iPSC-derived RPE cells from non-AMD human donors harboring the risk allele (402H CFH [homozygous; (H/H)] and observed that AKT2 phosphorylation is upregulated in *CFH* (H/H) cells compared to iPSC-derived RPE cells^21^ without the risk allele [controls; *CFH* (Y/Y)] (Fig. 1F). However, correction with the protective *CFH* I62V allele^18^ in *CFH* (H/H) RPE cells reduced the AKT2 phosphorylation levels (Fig. 1g), demonstrating a protective role of the *CFH* I62V variant in AKT2 signaling. Interestingly, *CFH* null (*CFH*^-/-^) iRPE (induced RPE) cells do not have elevated AKT2 (Fig. 1h). Furthermore, the increased AKT2 in the *CFH* (H/H) cells correlated with decreased Cathepsin D and L protein levels and activity (Fig. 2a-c), as also shown previously^20^. Notably, these lysosomal changes were not observed in human *CFH* null iRPE cells (Supplementary Fig. 3a, b). Collectively, these results link the *CFH* risk allele and elevated AKT2 to deficiencies in lysosomal biogenesis and compromised lysosomal function. However, the complete loss of CFH protein has no effect, raising the possibility that the 402H variant renders the CFH protein dysfunctional which mediates AKT2 induced lysosomal abnormalities. We also observed increased phosphorylation of both TFE3 and TFEB in RPE lysates from human AMD donors, which can be attributed to reduced nuclear translocation in the RPE of AMD patients; this was not found in age-matched controls (Fig. 2d). Accordingly, AMD patient eyes tend to show decreased nuclear TFEB immunostaining compared to non-AMD control eyes (Supplementary Fig. 4) and Cathepsin D and L protein levels and activities were decreased in human AMD donor RPE, compared to controls (Fig. 2e-g).

**Figure 2:**
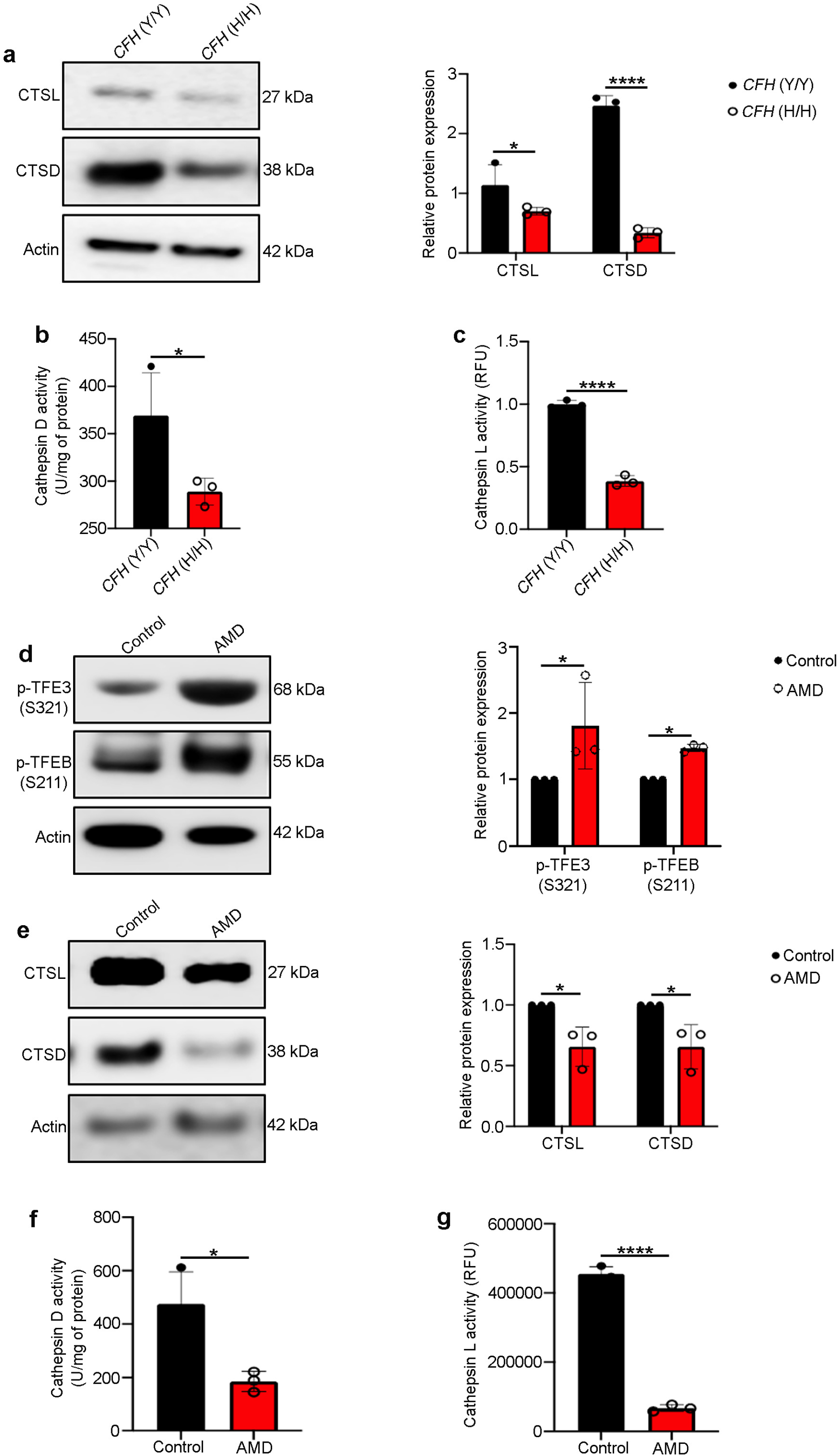
Akt2 upregulation in the RPE cells is associated with loss of lysosomal function in AMD. (**a**) Western blot and (**b**, **c**) colorimetric analysis revealed significant downregulation of (**a**) protein levels and (**b**, **c**) activities of lysosomal hydrolases CTSD and CTSL in iPSC-derived RPE cells from *CFH* Y402H risk allele [*CFH* (H/H)] containing donors, relative to controls. n=3. Western blot analysis showing (**d**) elevated levels of p-TFE3 (S321) and p-TFEB (S211) and (**e**) downregulation of CTSD and CTSL in RPE lysates from human AMD donors, compared to age-matched controls. n=3. RPE lysates from human AMD donors also showed significant downregulation of both (**f**) CTSD and (**g**) CTSL activities, compared to controls. n=3. All values are Mean ± S.D. ****P<0.0001, ***P<0.001, **P<0.01, *P<0.05.

We next set out to identify the mechanism by which increased AKT2 controls the nuclear translocation of TFEB/TFE3 and thereby lysosomal biogenesis. To this end, by performing a high throughput human protein-protein array^15^, we identified Sirtuin 5 (SIRT5) as the binding partner with the highest Z-score for AKT2 (Supplementary Table 1). This finding is of significant interest because sirtuins have been implicated in several metabolic and age-related diseases^22^. In particular, SIRT5 is an efficient deglutarylase, deacetylase, desuccinylase, and demalonylase of protein lysine residues, and thereby regulates both mitochondrial function and autophagy^23^. SIRT5 has also been shown to play a critical role in aging and has been identified as a possible therapeutic target for age-related diseases^22^. With computer modeling, we found that AKT2 (orange) could directly bind to SIRT5 (blue) (Fig. 3a).

**Figure 3:**
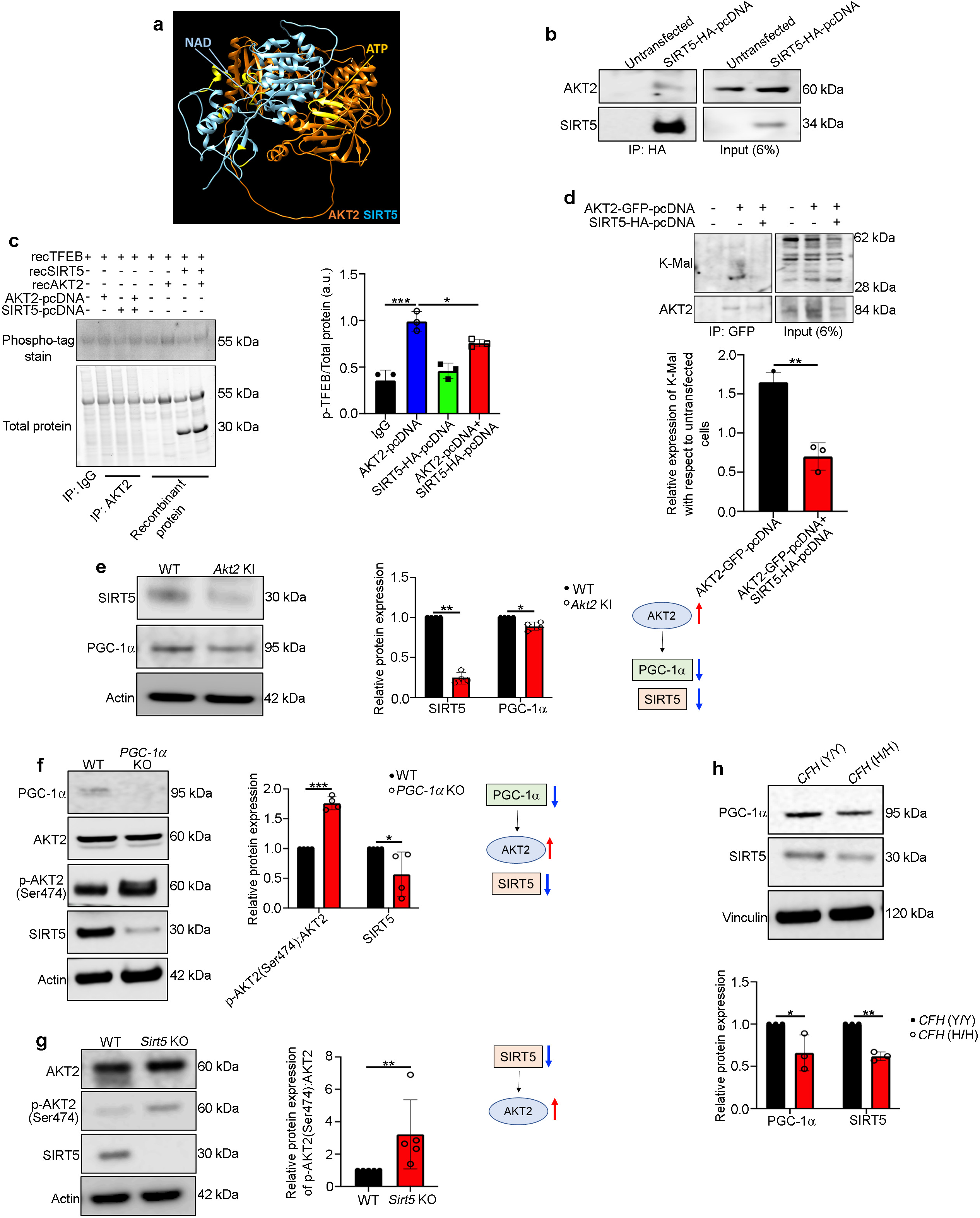
AKT2/SIRT5 signaling axis regulates TFEB activity. (**a**) A hypothetical heterodimer of human AKT2/SIRT5 suggests a potential protein-protein interaction between these proteins. The ribbon structures of AKT2 and SIRT5 are colored orange and cyan, respectively. Residues related to the AKT2 and SIRT5 active sites are shown in yellow. The model suggests that both proteins have active sites with potential exposure to substrates and ligands, indicating that the AKT2/SIRT5 complex is catalytically active. (**b**) Co-immunoprecipitation study showing that SIRT5 and AKT2 are binding partners in AKT2-pcDNA and SIRT5-HA-pcDNA overexpressing ARPE19 cells, compared to untransfected controls. n=3. (**c**) Pulldown of AKT2 with anti-AKT2 antibody and subsequent incubation of the pulldown complex with TFEB (recombinant; rec) *in vitro* to evaluate phosphorylation of TFEB using the phospho-tag gel staining, showed increased p-TFEB/total TFEB in AKT2 overexpressing ARPE19 cells, compared to control. This TFEB phosphorylation was reduced when cells were overexpressing both AKT2 and SIRT5, indicating SIRT5 can modulate AKT2 activity and subsequent TFEB phosphorylation. n=3. (**d**) Pulldown assay using anti-GFP magnetic beads from ARPE19 lysates overexpressing either GFP-AKT2 or GFP-AKT2 and SIRT5-HA, showed a decrease in lysine malonylation (K-Mal) of AKT2 pull down complex in presence of SIRT5 overexpression, compared to cells overexpressing only GFP-AKT2. n=3. Western blot analysis showing reduced expression of SIRT5 and PGC-1α in RPE cells from (**e**) *Akt2* KI, (**f**) *PGC-1α* KO, and (**g**) *Sirt5* KO mice, as well as (**h**) iPSC-derived RPE cells from *CFH* (H/H) donors, compared to controls. n=4 (mice) and n=3 (iPSC cells). All values are Mean ± S.D. **P<0.01, *P<0.05.

Co-immunoprecipitation studies in ARPE19 cells overexpressing AKT2-GFP and SIRT5-HA, demonstrated that SIRT5 binds to AKT2 (Fig. 3b). To determine if SIRT5 regulates AKT2 kinase activity *in vitro*, we incubated anti-GFP pulldown complexes from ARPE19 cells overexpressing AKT2-GFP or AKT2-GFP+SIRT5-HA with recombinant human TFEB and found that TFEB phosphorylation was reduced by AKT2 pulldown complexes from cells that also overexpressed SIRT5, when compared to cells overexpressing only AKT2 (Fig. 3c).

Among the post translational modifications targeted by SIRT5, malonylation is known to be prominent in both cytosol and mitochondria^22, 23^. In contrast, succinylation, acetylation and glutarylation are prominent in mitochondria only^22, 23^. Since, SIRT5 and AKT2 both localize to cytosol and mitochondria^6, 15, 16, 22, 23^, we therefore performed anti-GFP pulldown experiments from AKT2-GFP and SIRT5-HA+AKT2-GFP overexpressing ARPE19 cells to assess lysine malonylation (K-Mal) of AKT2-GFP complex proteins. K-Mal formation was significantly reduced in cells overexpressing both SIRT5 and AKT2 relative to cells overexpressing only AKT2 (Fig. 3d). These results indicate that SIRT5 can reduce the malonylation of AKT2 complexes, which in turn, regulates AKT2 activity. We next assessed the crosstalk between SIRT5 and AKT2 in RPE cells. Since AKT2 can inhibit peroxisome proliferator-activated receptor-coactivator 1 alpha (PGC-1α)^24^, and PGC-1α upregulates SIRT5^25^, we reasoned that in the RPE of *Akt2* KI mice, which overexpress AKT2 in the RPE, SIRT5 abundance would be mediated by the AKT2/PGC-1α axis. We observed that both PGC-1α and SIRT5 expression were decreased in the RPE of *Akt2* KI mice compared to WT mice (Fig. 3e). Conversely, *Akt2* cKO RPE cells demonstrated an increase in both proteins relative to floxed controls (Supplementary Fig. 5). Furthermore, SIRT5 was decreased and p-AKT2 (Ser474) was increased in RPE lysates from both *PGC-1α*^-/-^ and *Sirt5*^-/-^ mice^26, 27^ (Fig. 3f, g). Importantly, in *CFH* (H/H) iPSC-RPE cells, which have high AKT2 signaling, both SIRT5 and PGC-1α were decreased as compared to control *CFH* (Y/Y) cells (Fig. 3h). Collectively, these results suggest that AKT2 regulates SIRT5 expression through PGC-1α, whereas SIRT5 can regulate AKT2 activity and subsequently phosphorylate AKT2 targets, including TFEB.

Since PGC-1α can regulate mitochondrial biogenesis^25, 26^, we next explored mitochondrial function in ARPE19 cells overexpressing either AKT2 or AKT2+SIRT5 by measuring the oxygen consumption rate (OCR) and extracellular acidification rate (ECAR) using the Seahorse assay^28^. While both maximal respiration and ATP-linked respiration were impaired in AKT2 overexpressing cells, they were rescued by the additional SIRT5 overexpression (Supplementary Fig. 6a-c). These data suggest that a functional connection between lysosomes and mitochondria is essential for maintaining RPE homeostasis. While dysfunction of either organelle has been shown to play a pivotal role in AMD pathobiology, these results show that the lysosome-mitochondria crosstalk is also important for maintaining RPE health.

Lysosomal dysfunction contributes to RPE degeneration and the formation of extracellular deposits in Bruch’s membrane called drusen, two hallmarks of early AMD^1, 3, 4^. The pathogenesis of drusen remains unclear. We set out to determine if malfunction of lysosome-mediated physiological processes, such as autophagy, could drive the extracellular waste accumulation that contributes to drusen biogenesis. Secretory autophagy is a ‘non-canonical’ phenomenon in which autophagosome cargo is released from the cell through the plasma membrane by a lysosome-independent pathway^29^. Synaptotagmin Like 1 (SYTL1) protein, which is a critical regulator of secretory autophagy^30^, was another binding partner of AKT2 in our high throughput human protein-protein interaction study (Supplementary Table 1). Therefore, we determined whether secretory autophagy is activated by increased AKT2 signaling. Using a pulldown assay, we confirmed that AKT2 associates with SYTL1 (Supplementary Fig. 7a) and forms a complex also containing tripartite motif-containing protein 16 (TRIM16)^29^ and synaptosome-associated protein 23 (SNAP23)^29^. This complex is needed for efficient release of cellular cargo from cells during secretory autophagy (Supplementary Fig. 7b). Intriguingly, formation of this secretory autophagy complex was inhibited by overexpressing SIRT5 (Supplementary Fig. 7b), likely due to the regulation of AKT2 by SIRT5.

AKT2 overexpression in ARPE19 cells led to upregulation of other secretory autophagy mediators including FK-506-binding protein 51 (FKBP51)^31^ and SNAP23, which was not seen in cells overexpressing AKT2+SIRT5 or a mutant form of AKT2 (K14A/R25E)^32^ that has reduced kinase activity (Fig. 4a). Moreover, we observed that GFP-LC3 positive autophagosomes do not fuse with lysosomes in AKT2 overexpressing ARPE19 cells, which indicates that autophagosome clearance is impaired (Fig. 4b). Total internal reflection fluorescence (TIRF)^33^ microscopy studies confirmed that the autophagosomes fused with the plasma membrane, but not with lysosomes (Supplementary Movies 1,2 and Supplementary Fig. 7c).

**Figure 4:**
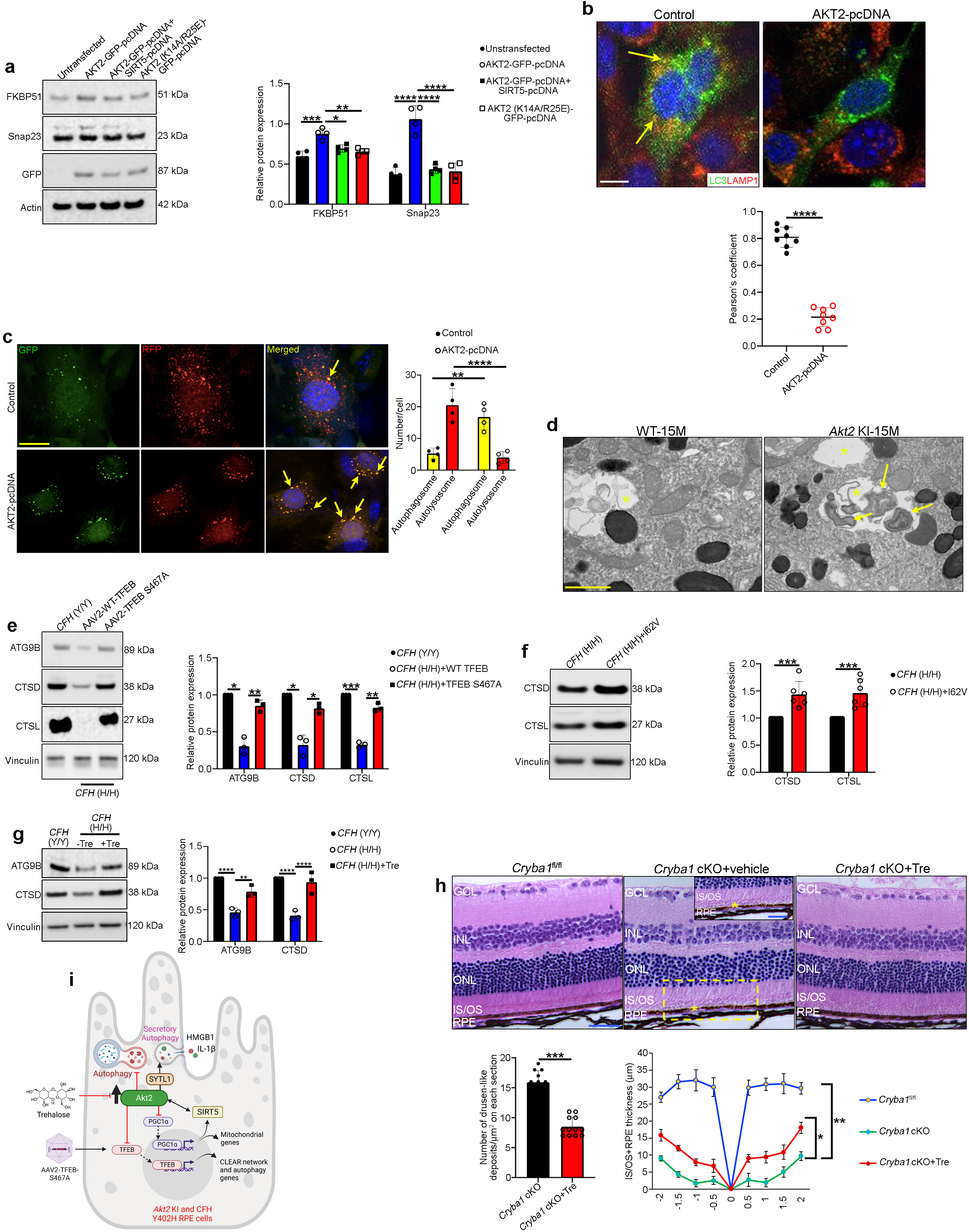
Akt2 upregulation triggers secretory autophagy in RPE cells; targeting TFEB independent of mTOR can rejuvenate lysosomal function. (**a**) Western blot showing elevated levels of FKBP51 and Snap23 in AKT2 overexpressing ARPE19 cells, which were reduced upon simultaneous upregulation of SIRT5 or overexpression of an AKT2 inactive mutant (K14A/R25E). n=4. (**b**) Immunofluorescence assay showing association of LC3-positive autophagosomes (green) with lysosomes (Lamp1-positive; red) in control ARPE19 cells (arrows in **b**), which was significantly reduced upon AKT2 overexpression. n=8. (**c**) ARPE19 cells were transfected with AKT2 construct for 48 h or left untreated (control), followed by an overnight infection with an Adenovirus-GFP-RFP-LC3B construct to label the autophagosomes (yellow) and autolysosomes (red). The number of autolysosomes (red puncta) was significantly decreased in AKT2 overexpressing cells (arrows in **c**) when compared to controls, suggesting decline in autophagy flux. n=4. (**d**) Transmission electron micrographs showing double membranous autophagosomes in the RPE cells of 15-month-old *Akt2* KI mice, but not in age-matched WT. n=5. Western blot analysis showing that in iPSC-derived RPE cells from *CFH* (H/H) risk allele-containing donors, overexpression of (**e**) mutant TFEB S467A construct (10^6^ vg/ml for 48 h) or (**f**) treatment with trehalose (100 mM for 20 h) or (**g**) correction with *CFH* I62V mutation rescued the levels of the lysosomal hydrolases Cathepsin D, Cathepsin L and the autophagosome mediator ATG9B, compared to untreated *CFH* (H/H) cells. n=3 (TFEB S467A and trehalose groups), n=6 (*CFH* I62V rescue). (**h**) Hematoxylin-eosin stained sections and spider plot showing decrease in IS/OS+RPE thickness and patchy RPE layer with accumulation of drusen-like deposits (asterisks in **h** and inset) in *Cryba1* cKO retina (vehicle control), with rescue by trehalose treatment. n=5. (**i**) Created with BioRender.com. Cartoon showing that AKT2 upregulation in RPE cells from *Akt2* KI mice and *CFH* (H/H) iPSC-derived cells trigger deregulation of SIRT5/PGC-1α dependent lysosomal function. Trehalose and S467A TFEB mutant construct rescued the lysosomal abnormalities and delayed the progression of early RPE changes in a mouse model. All values are Mean ± S.D. ****P<0.0001, ***P<0.001, **P<0.01, *P<0.05.

The conditioned medium from 10-month-old *Akt2* KI RPE explants had significantly increased major secretory autophagy cargo^29^, such as interleukin-1 beta (IL-1β) and high mobility group box 1 (HMGB1), compared to WT RPE cells (Supplementary Fig. 7d). A similar increase was observed in the levels of these cargo proteins in the conditioned medium from *CFH* (H/H) iPSC RPE cells (Supplementary Fig. 7e) . Since secretory autophagy and canonical degradative autophagy are integrated and highly regulated processes that share similar molecular mechanisms^29^, we next showed that canonical autophagy was reduced in *Akt2* KI RPE cells (Supplementary Fig. 8a-c). RNAseq analysis of *Akt2* KI RPE cells identified a cluster of downregulated genes involved in autophagosome formation^34^, including autophagy related 9B (*Atg9b*) and the Unc-51-like kinase 1 (*Ulk1*) (Supplementary Fig. 8a). Their reduced protein abundance as well as accumulation of the autophagosome marker, sequestosome 1 (p62/SQSTM1)^14^, was confirmed by western blots (Supplementary Fig. 8b). Genes involved in autophagosome formation are transcriptionally regulated by TFEB. Using chromatin immunoprecipitation (ChIP)^12^, we found that TFEB fails to bind to the promoter of *Atg9b* in *Akt2* KI RPE cells (Supplementary Fig. 8c). These abnormalities correlated with a decline in autophagy flux^14, 35^ in RPE cells overexpressing AKT2 (Fig. 4c and Supplementary Fig. 8d). Moreover, double membranous autophagosomes^35^ accumulated in the RPE of *Akt2* KI mice by 15 months of age, but were not seen in age-matched WT (Fig. 4d). Collectively, these results suggest that AKT2 upregulation in the RPE induces alterations in secretory autophagy that promote the release of material that contributes to drusen formation.

Based on our findings, we speculated that targeting the AKT2/TFEB pathway in mouse AMD models, as well as in *CFH* (H/H) iPSC-derived RPE cells, could prevent early changes observed in AMD pathogenesis. We therefore targeted AKT2-dependent TFEB signaling in *CFH* (H/H) iPSC-derived RPE cells with (1) AAV2-TFEB-S467A, a mutation in the AKT-target residue of TFEB that induces constitutive nuclear localization^6^, (2) treatment with the disaccharide trehalose, which activates TFEB independent of mTOR signaling^36^, or (3) correction with the protective *CFH* I62V mutation. These treatments rescued the levels of lysosomal and autophagy mediators (Fig. 4e-g). Further, trehalose rescued CLEAR gene expression and autophagy flux in *Cryba1* KO RPE explants^35^ (Supplementary Fig. 9a-c) without influencing the abnormal mTORC1 activation (Supplementary Fig. 9d). This finding suggests that trehalose can rejuvenate lysosomal function independent of mTOR signaling. These findings encouraged us to investigate whether trehalose could prevent the lysosomal abnormalities observed in our mouse models^14, 16^ that develop an AMD-like phenotype. Trehalose administered^37^ to 6-month-old *Cryba1* cKO or *Akt2* KI mice for 3 consecutive months rescued CLEAR network gene expression including autophagy mediators, and restored lysosomal function in the RPE (Supplementary Fig. 10a-e). Moreover, while Bruch’s membrane basal deposits are typically seen in *Cryba1* cKO mice by 9 months of age^14, 15^, they were not visible in trehalose-treated mice (Fig. 4h). Thus, targeting TFEB, rescued autophagy and lysosomal biogenesis in both *CFH* (H/H) iPSC-derived RPE cells and mouse models with lysosomal deficits (Fig. 4i). Importantly, the rescue of lysosomal dysfunction prevented basal laminar deposit formation in our mouse AMD model.

Since vision loss develops late in AMD^1^, patients often present to their eyecare doctor only after the underlying pathology has progressed significantly. While it is conceptually feasible to correct the Y402H *CFH* risk variant by CRISPR/Cas9 editing in human patients as a future therapy^38^, this may not prove to be ideal because of the nature and stage of the disease at diagnosis. While mTORC1 has been a popular therapeutic target due to its regulation of TFEB/TFE3 nuclear localization and lysosomal activity, these studies have been unsuccessful because of intolerable side effects^39^. Given the lysosomal defects in atrophic AMD^3, 4^, especially in patients who harbor the Y402H *CFH* risk variant, and important interactions between lysosomes and mitochondria, our results provide an alternative strategy targeting specific relevant downstream pathways. Activating the lysosome/autophagy pathway, and its communication with mitochondria, represents a novel potential therapeutic target that could rejuvenate both organelles that become impaired in atrophic AMD.

## Methods

### Antibodies

The primary antibodies AKT2 (3063S), p-AKT2 (8599S), SIRT5 (8782S), CTSD (69854S), ULK1 (8054T), FKBP51 (8245S), p-TFEB (37681S), K-Malonyl (14942S) and GFP (2555S) were purchased from Cell Signaling Technology. CTSL (NB100-1775), p62/SQSTM1 (NBP1-42821) and PGC-1α (NBP1-04676) were purchased from Novus Biologicals. TFE3 (14480-1-AP), ATG9B (PA5-20998), Ezrin (PA518541), Trim16 (PA5-110515), and Snap23 (PA1-738) were purchased from Thermo Fisher. Loading controls were vinculin (ab129002, Abcam) and beta-Actin (4970S, Cell Signaling). TFEB (A303-673A) and SYTL1 (A305-648A-T) antibodies were purchased from Bethyl Laboratories. The p-TFE3 antibody was a gift from Dr. Rosa Puertollano’s laboratory. The secondary antibodies used in this study were purchased from KPL: anti-rabbit (074–1506) and anti-mouse (074–1806).

### Animals and trehalose treatment

All animal studies were conducted in accordance with the Guide for the Care and Use of Animals (National Academy Press) and were approved by the University of Pittsburgh Animal Care and Use Committee. Both male and female RPE-specific *Akt2* KI, *Akt2* cKO, and βA3/A1-crystallin conditional (*Cryba1* cKO), as well as βA3/A1-crystallin complete knockout (*Cryba1* KO), *PGC-1α* KO, and *Sirt5* KO mice were generated as previously described^14–16, 26, 27^. All mice that were used in this study were negative for RD8 mutation^14^. At 6 months of age, the *Cryba1* cKO and *Akt2* KI mice were injected with trehalose (T9449-100G, Sigma Aldrich) intraperitoneally at 1 mg/g/day for 2 days and then given 2% trehalose (w/v) in drinking water *ad libitum*^37^ for the entire experimental duration of 3 months.

### iPSC-derived cells culture

Human iPSC-derived RPE cells were obtained from genetically screened human donors as explained previously^40–42^. Both the control, *CFH* (Y/Y)- and the risk allele-*CFH* (H/H) containing donor cells were differentiated and maintained as described previously^41^. *CFH* (H/H) cells with the rescue *CFH* I62V mutation and *CFH* null cells were also generated and maintained as described earlier. The iPSC lines were cultured on vitronectin-coated plates in E8 complete medium (A1517001, Thermo Fisher)^41^. The recombinant vitronectin (A14701SA, Thermo Fisher) was diluted in DPBS at the concentration of 1:200 to get the final concentration of 2.5 mg/ml. The cells were passaged every 4-5 days at 70-80% confluency^41^ and were fed every 24 hours. The differentiated cells were confirmed for RPE markers and were subjected to downstream experiments.

### Human AMD donors

The human AMD donor sections were purchased from Lion’s Gift of Sight, Minnesota. The control (n=5) and AMD (n=3) donors were all Caucasians with an average age of 76 ± 5 years. The control donors did not have any eye diseases. The AMD donors were staged according to the Minnesota Grading System (MGS) and all the donors were classified as MGS3, as explained earlier^12^.

### RPE flatmount and immunostaining

After euthanizing WT and *Akt2* KI mice, the eyes were immediately enucleated using angled forceps. Eyes were briefly washed in 1X PBS at RT and dabbed on Kimwipes to remove PBS. The eyes were then introduced into tubes containing 4% paraformaldehyde (PFA). After 2 h of fixation eyes were cleaned by trimming away the extra-ocular tissue and debris under the dissecting microscope, leaving the optic nerve intact^15, 17^. The cleaned eyes were again added to the tubes containing PFA, and left for another 3 h at RT. The fixed eyes were removed from PFA, added to 30% sucrose and for 10-12 h at 4°C. The eye balls were then dissected under the dissecting microscope for RPE flatmount preparation as described previously^15, 17^. To eliminate autofluorescence from the melanin in the RPE flatmount, the RPE cups were treated with 1 mg/ml solution of sodium borohydride for 30 mins. This was followed by permeabilization and blocking steps as previously described^15, 17^. The flatmounts were incubated with with ZO-1 (1:200) (13663S, Cell Signaling Technology) in primary antibody buffer (1% Tween 20 + 0.5% BSA in PBS) and incubated, overnight at 4°C. The primary antibody was removed, and the RPE flatmounts were washed with PBS+1% Tween 20. The flatmounts were then incubated with Anti-Rabbit Alexa fluor 555 secondary antibody (1:500) (A31572, Invitrogen) prepared in secondary antibody buffer (0.1% Tween 20 + 0.5% BSA in PBS), along with Hoechst (1:2000) (62249, Thermo Fisher), for nuclear staining and incubated in the dark for 2 h at RT^15, 17^. The slides were then washed with 1X PBS, 5 times, for 15 minutes each and then mounted with DAKO mounting media before imaging using an Olympus IX81 confocal microscope^15, 17^.

### RPE explant culture

Immediately after euthanization of 3 month old WT or *Cryba1* KO or *Akt2* KI mice, the eyes were enucleated, and anterior segments were removed. The posterior eye cups were then cut into four petals and the neural retina was carefully removed^17, 35^. The harvested RPE-choroid-sclera (RCS) complex was placed onto PVDF membranes after flattening them by making several relaxing cuts. The explants were then cultured face up in complete media as previously described^17, 35^. Treatment with trehalose was done on *Akt2* KI or *Cryba1* KO RPE explants at a dose of 100 mM for 20 h as explained previously^6, 17, 35^.

### Mouse retinal cryosectioning

Eyes from freshly euthanized WT and *Akt2* KI mice were fixed following the same procedure for RPE flatmount preparations^15, 17^. The cryosectioning of the eye globes and subsequent permeabilization and blocking of the sections was performed as explained previously^15, 17^. The slides were incubated with primary antibody cocktail containing either rhodopsin (ab98887, Abcam) or EBP50 (PA1090, Thermo Fisher), in primary antibody buffer (1% Tween 20 + 0.5% BSA in PBS) at a dilution of 1:100 and incubated in a moist chamber overnight at 4°C. The slides were washed with PBS+1% Tween 20 three times. Secondary antibody cocktail Anti-mouse Alexa fluor 647 (1:200) and Anti-Rabbit Alexa fluor 555 (1:200) was prepared in secondary antibody buffer (0.1% Tween 20 + 0.5% BSA in PBS) also containing Hoechst (1:2000) and Alexa fluor 488-phallodin (A12379, Thermo Fisher) (1:1000) and was added to each slide covering the sections completely^15, 17^. The slides were washed with 1X PBS, 5 times, for 15 min each and then mounted with a coverslip using DAKO mounting medium until imaged with an Olympus IX81 confocal microscope^15, 17^. The length of phalloidin-positive apical microvilli on RPE cells was quantified using ImageJ software^15, 17^.

### Immunostaining and quantification of TFEB nuclear translocation using paraffin-embedded human retinal sections

Human control and AMD donor paraffin-embedded eye globes were subjected to dewaxing using three washes of fresh xylene and rehydration using a gradient of ethanol followed by antigen retrieval as explained previously^15, 17^. The slides were then washed with PBS/0.1% Tween 20, twice for 10 min each and then subjected to permeabilization and blocking steps as previously demonstrated^15, 17^. The slides were incubated with TFEB primary antibody (A303-673A, Bethyl Laboratories) in primary antibody buffer (1% Tween 20 + 0.5% BSA in PBS) at a dilution of 1:100 and incubated in a moist chamber overnight at 4°C. The slides were washed with wash buffer three times. A secondary antibody cocktail was prepared containing Anti-Rabbit Alexa fluor 647 (1:200) in secondary antibody buffer (0.1% Tween 20 + 0.5% BSA in PBS) also containing Hoechst (1:2000), and was added to each slide covering the sections completely and incubated in dark for 2 h at RT. Post-staining the slides were washed with 1X PBS, 5 times, for 15 min each and then mounted with a coverslip using an DAKO mounting media until imaging in a confocal microscope^15, 17^. Slides were imaged using VS200 Olympus Slide Scanner with 40x objective (UPlanXApo NA 0.95) in spectral ranges: DAPI (ex. 378/52, em. 432/36), Cy3 (ex. 554/23, em. 595/31), Cy5 (ex. 635/18, em. 680/42) and Cy7 (ex. 735/28, em. 809/81) with ORCA Fusion BT digital CMOS camera. Cy7 channel was used to detect RPE layer by thresholding followed by morphological operations. Individual nuclei were detected using Cellpose^43^ generalist segmentation algorithm (model ‘cyto’) in DAPI channel. TFEB specific signal was calculated as a difference between Cy5 and Cy3 channels. Individual foci were detected using Difference of Gaussian algorithm (scikit-image^44^). A nucleus was considered positive if more than two foci were detected within its area.

### Transmission electron microscopy

The WT and *Akt2* KI mouse eyes were immersed fixed in cold 2.5% glutaraldehyde (25% glutaraldehyde stock EM grade, Polysciences, Warrington, PA) and 2% paraformaldehyde (Fisher Scientific, Pittsburgh, PA) in 0.01 M PBS (sodium chloride, potassium chloride, sodium phosphate dibasic, potassium phosphate monobasic, Fisher Scientific, Pittsburgh, PA), pH 7.3. The sclera, choroid, and retina were separated and quartered after the optic nerve was removed^35^. The sample was rinsed 3x in PBS, post-fixed in 1% osmium tetroxide (Electron Microscopy Sciences) with 1% potassium ferricyanide (Fisher Scientific) for one hour, dehydrated by putting the samples in a series of ethanol (30% - 90%), and propylene oxide (Electron Microscopy Sciences). Next, the samples were properly oriented and embedded in Poly/Bed® 812 (Dodecenyl Succinic Anhydride, Nadic Methyl Anhydride, Poly/Bed 812 Resin and Dimethylaminomethyl, Polysciences). Thin (300 nm) cross sections of the area adjacent to the optic nerve were cut on a Leica Reichart Ultracut (Leica Microsystems), stained with 0.5% Toluidine Blue in 1% sodium borate (Toluidine Blue O and Sodium Borate, Fisher) and examined under the light microscope. Ultrathin sections of the same area (65 nm) were stained with 2% uranyl acetate (Electron Microscopy Sciences) and Reynold’s lead citrate (Lead Nitrate, Sodium Citrate and Sodium Hydroxide, Fisher) and examined on JEOL 1400 Flash transmission electron microscope with AMT Biosprint 12, 12mp camera (Advanced Microscopy Techniques).

### RNAseq analysis and bioinformatics

The RNAseq analysis was performed from RPE cells harvested from WT and *Akt2* KI mice at 4 months of age as a paid service from Novogene Inc. The bioinformatics analysis was performed to identify differentially changing autophagy genes using previously published gene sets. Average expression values of the particular genes in each sample were then calculated by the function “AverageExpression” in the Seurat package^35^.

### Chromatin Immunoprecipitation

Chromatin Immunoprecipitation (ChIP) was performed as previously described^12^. RPE cells were harvested from WT and *Akt2* KI mice and ChIP was performed using a commercially available kit (17-295, EMD Millipore). The following primers for the Atg9b promoter and negative control were used Atg9B_CHIP_F1: CACATGCTAGAGCCCTCTGATA, Atg9B_CHIP_R1: GGCTCAAGAGCTATTGGGATT, Atg9B_CHIP_F2: GGCATGTGGCCTTAAATCAT, Atg9B_CHIP_R2 GGCTCTAGCATGTGCAGACA, Atg9B_CHIP_NEG_F: TGGCCATATCTGGGAGTCAA, Atg9B_CHIP_NEG_R: CCAGCCAGGCTAGTGTTCTC.

### Co-immunoprecipitation

ARPE19 cells were either transfected with EGFP-AKT2 or HA-SIRT5 or EGFP-AKT2 (K14A/R25E) (86593; 24483; 86595, Addgene) constructs using a Lipofectamine transfection kit (L3000008, Thermo Fisher)^14, 15^. After 48 h the cells were lysed and a pulldown assay was performed from cell lysates using either anti-HA (88836, Thermo Fisher) or anti-GFP (gtd-100, ChromoTek) magnetic beads followed by western blotting for respective proteins of interest.

### Human high-throughput protein-protein interaction

The evaluation of AKT2 binding partners was performed as a paid service from CDI NextGen Proteomics, Baltimore, MD, USA, as previously described^15, 17, 35^. Briefly, recombinant AKT2 was used for this study using the HuProtTM v3.1 human proteome array and the sample was placed on array plates at a concentration of 1 μg/mL. The results were analyzed using GenePix software as described previously and represented as the significance of the probe binding signal difference from random noise (Z-Score). Only protein interactions with a Z-score above 6 were considered^15, 17, 35^.

### Molecular modelling

Models of molecular structure for human RAC-beta serine/threonine-protein kinase (AKT2) and NAD-dependent protein deacylase sirtuin-5 (SIRT5) were obtained from the AlphaFold Protein Structure Database (ebi.ac.uk), models AF-P31751-F1-model_v4.pdb and AF-Q9NXA8-F1-model_v4.pdb, respectively. Molecular modeling of the possible interaction between AKT2 and SIRT5 was done using the ‘movement’ tool in UCSF CHIMERA, VER. 1.17.1.

### In vitro AKT2 activity measurement

ARPE19 cells were overexpressed with EGFP-AKT2 with or without HA-SIRT5 construct. After 48 h, the cells from all the experimental groups were lysed in CHAPS lysis buffer, followed by pulldown with anti-GFP magnetic beads. The AKT2 complex was incubated with recombinant TFEB (TP760282, Origene) for 15 minutes at 37°C in the presence of 10 nm ATP (A1852-1VL, Sigma Aldrich). The mixture was denatured using LDS sample buffer (NP0007, Invitrogen) containing β-marcaptoethanol and then SDS-PAGE was run. Phospho-protein staining was performed using a commercially available kit (P005A, ABP Biosciences), to evaluate the levels of TFEB phosphorylation. The phospho-protein band intensity in each experimental group was calculated by ImageJ software and graphical representation was performed as the ratio of the phospho-protein to the total protein band intensity for each sample^35^.

### Quantitative polymerase chain reaction (qPCR)

The expression of different genes was evaluated by qPCR as described previously^35^, using a commercially available cDNA preparation kit (11754-050, Invitrogen) and Taqman probe-based qPCR master mix (4369016, Thermo Fisher). The commercially available Taqman probes (Thermo Fisher) used for this assay were; *Ctsd* (Mm00515586), *Ctsb* (Mm01310506), *Lamp1* (Mm00495262) and Atp6v0a1 (Mm00444210).

### Western blot

Western blotting was performed using previously published methods from our laboratory^15, 17, 35^. The cell and tissue lysates were prepared in 1X RIPA buffer (20–188, EMD Millipore) containing 0.1% of protease inhibitor cocktail (I3786, Sigma Aldrich) and 0.1% phosphatase inhibitor cocktail (P0044-5ML, Sigma Aldrich). Densitometry was performed to estimate the protein expression relative to the loading control (Actin) using ImageJ software (National Institute of Health)^15, 17, 35^.

### Enzyme-linked immunosorbent assay (ELISA)

To evaluate the levels of IL-1β and HMGB1 in spent medium from WT and *Akt2* KI RPE explant cultures^15^, as well as in *CFH* (Y/Y) and *CFH* (H/H) iPSC-derived RPE cell cultures, commercially available kits were purchased and the assays done following the manufacturer’s protocol: IL-1β (Mouse: BMS6002; Human: BMS224INST, Thermo Fisher) and HMGB1 (Mouse: NBP2-62767; Human: NBP2-62766, Novus Biologicals).

### Estimation of cathepsin D and L activities

The estimation of cathepsin D (ab65302, Abcam) and L (ab65306, Abcam) activities from RPE lysates from WT or *Akt2* KI mice or from iPSC-derived RPE cells was performed by using commercially available kits.

### Generation of TFEB KO MEFs

*Tfeb* KO stable cell line was generated on the Trf2F/F; Rosa-CreER MEFs background^45^. The cells were infected with LentiCRISPR.v2 puro construct carrying the following guide sequences: Tfeb-3F: caccgCAGCCCGATGCGTGACGCCA and Tfeb-3R: aaacCTGGCGTCACGCATCGGGCTG which targets exon 1 of mouse *Tfeb* gene. After infection, cells were treated with puromycin for selection and single clones were isolated to get the *Tfeb* KO MEF cell clones for further propagation.

### Seahorse

ARPE19 cells were plated at 40,000 cells per well on a Seahorse XF platform compatible 96-well plate pre-coated with poly-D-lysine (Sigma Aldrich, USA) and allowed to adhere and grow for 24 h^28^. The cells were transfected with either EGFP-AKT2 or EGFP-AKT2+ HA-SIRT5 constructs or left untransfected and after 48 hours were subjected to the Mitostress assay kit (Cat# 103015-100) from Agilent, USA^28^.

### Hematoxylin-eosin staining

Eyes from *Cryba1*-floxed and trehalose or vehicle (water) treated *Cryba1* cKO mice were fixed in 2.5% glutaraldehyde followed by formalin, and then subjected to ethanol gradation and dehydrated followed by embedding in methyl methacrylate^15^. Sections (1 μm) were cut and stained with hematoxylin and eosin and observed under a light microscope as described previously. The RPE+IS/OS thickness and the quantification of drusen-like deposits were quantified as explained previously^15^.

### Fundus imaging

Fundus photographs were obtained from WT and *Akt2* KI mice using the Micron IV Laser Scanning Ophthalmoscope (Phoenix Research Lab, Inc) as previously described^14^.

### Electroretinography

WT and *Akt2* KI mice at 4 and 10 months of age were dark adapted for 20 h and then were anesthetized with ketamine (50 mg/kg body weight) and xylazine (10 mg/kg body weight) and subjected to electroretinography to evaluate retinal function by estimating the scotopic a-, b- and c-wave responses using the Celeris Diognosys System, USA^16^. Responses were measured at three different light intensities (0.01, 0.1 and 1 cd*s/m2)^16^.

### Autophagy flux

Autophagy flux on RPE explants from WT and *Akt2* KI mice as well as *Cryba1* KO (+/-trehalose; 100 mM for 24 h) mice was performed as described previously^35^. Briefly, the explants were treated with Bafilomycin A1 (BafA1; 1μm) or chloroquine (ChQ; 50 μM) or left untreated for 6 h^14, 35^. Western blotting was performed to evaluate the levels of LC3-I and LC3-II. Autophagy flux was estimated by calculating the ratio of LC3-II in BafA1 or ChQ-treated cells with respect to untreated cells^35^.

### Immunofluorescence analysis of autophagosome/autolysosome formation and lysosome/autophagosome fusion

To evaluate the levels of autophagosome and autolysosome formation upon AKT2 overexpression in RPE cells, the ARPE19 cells were transfected with either an AKT2-pcDNA construct (86623, Addgene) or left untreated for 48 h and then infected for 12 h with 10^3^ vg/ml of an Adenovirus-GFP-RFP-LC3 construct (Vector Biolabs, 2001)^35^. The cells were fixed with 2% PFA for 30 min at 4°C and the nuclei were stained with Hoechst at 1 mg/100 ml dH_2_O. Confocal images were acquired on the Zeiss LSM 710 platform (Switzerland). The data were analyzed using ImageJ software as previously described^35^ and represented as puncta number for autophagosomes (yellow) and autolysosomes (red) in control and AKT2 overexpressing cells. Lysosome/autophagosome fusion was estimated in ARPE19 cells overexpressing only GFP-LC3 (24920, Addgene) or both GFP-LC3 and AKT2 constructs. After 48 h the cells were fixed (as mentioned above) and then immunostained with Lamp1 antibody (ab24170, Abcam), followed by secondary antibody staining (Goat Anti-rat Alexa fluor 455; 1:300) as explained previously^14, 15^. Analysis of colocalization between LC3 (green) and Lamp1 (red) was performed using the JACoP plugin of ImageJ software to evaluate Pearson’s co-efficient and the level of colocalization between the two proteins was estimated as previously described^35^.

### Total internal reflection fluorescence (TIRF) microscopy

TIRF imaging was performed using an iLas2 Ring TIRF illuminator on a Nikon Ti inverted microscope equipped with a 1.49 N.A. 100x Nikon TIRF objective and 1.5x tube lens with 0.07 effective pixel size^33^. Images were acquired using a Photometrics Prime 95B sCMOS camera. Images of the ARPE19 cells with or without AKT2 overexpression for 48 h followed by infection with 10^3^ vg/ml of Adenovirus-GFP-RFP-LC3 construct for 12 h were collected in serum-free starvation medium for 2 hours^35^. Images were analyzed using NIS Elements spot detection in order to identify punctate structures at the membrane. The number of spots were counted over time and each field of view (FOV) was normalized to the average number of spots in the FOV to compare relative changes between all analyzed FOVs. The quantification was performed using Graph Pad Prism 8 software to ascertain the difference in between the two groups across all the time points using linear regression plots.

### Statistical analysis

All analyses were performed using the GraphPad 8.0 software. Student’s t-test and one-way ANOVA followed by Tukey post hoc test were used to measure the statistical differences between groups^15, 35^. The significance was set at P < 0.05. The analyses were performed on triplicate technical replicates^15, 35^. All the values are presented as mean ± standard deviation (SD)^15, 35^.

## Acknowledgements

This work was supported by NIH R01 EY031594 (to DS and JTH), NIH R01 EY032516 (to DS), Edward N. and Della L. Thome Memorial Foundation Awards Program in Age-Related Macular Degeneration Research (to DS), BrightFocus Foundation Postdoctoral Fellowship on Macular Degeneration (to SG), start-up funds to DS from Ophthalmology, University of Pittsburgh, the Jennifer Salvitti Davis, M.D. Chair Professorship in Ophthalmology (DS), The Robert Bond Welch Professorship (JTH), P30 core award EY08098 from the National Eye Institute, NIH (to the University of Pittsburgh Department of Ophthalmology), and unrestricted funds from The Research to Prevent Blindness Inc., NY (to the University of Pittsburgh Department of Ophthalmology and the Wilmer Eye Institute) and the Academy of Finland grant 333302 (to KK). The authors would also like to acknowledge the bioinformatics and imaging core facilities within the Department of Pediatrics, University of Pittsburgh School of Medicine and Kaarniranta AMD lab staff for *PGC-1α* KO mice generation and sample isolation. This project has also been made possible in part by grant number 2020-225716 from the Chan Zuckerberg Initiative DAF, an advised fund of Silicon Valley Community Foundation.

## Data Availability Statement

All data generated or analysed during this study are included in this published article (and its Supplementary Information files). The RNAseq data for the genes depicted in Supplementary Fig. 8 will be uploaded in the NCBI GEO database after acceptance. Figure 4i was created with BioRender.com.

## Author Contributions

DS designed and conceptualized the study. J.T.H provided input about the genetic aspects of the study. S.G., S.B., V.K., M.N., M.Y. performed the experiments. R.S., D.B. K.B. generated the iPSC-derived RPE cells lines and the *CFH* null and isogenic control cells (iRPE). K.M.K. performed AI-based quantification of immunofluorescence data. C.T.W and S.C.W. performed and analysed TIRF experiment results. J.F. performed the TEM imaging. C.Y.W. and T.F. generated the *Tfeb* KO and *Tfeb*-flox MEF line. Y.S. performed the computer modelling. D.R. performed bioinformatics analysis. E.S.G. and K.K provided the *Sirt5* KO and *PGC-1α* KO mice. S.G., D.S., R.P., S.H., A.S., M.F.B., J.A.S., J.T.H. and J.S.Z. analysed the data. S.G., D.S., S.H., A.S., J.T.H. and J.S.Z wrote the paper.

