## Supplementary Information for "The AKT2/SIRT5/TFEB pathway as a potential therapeutic target in atrophic AMD"

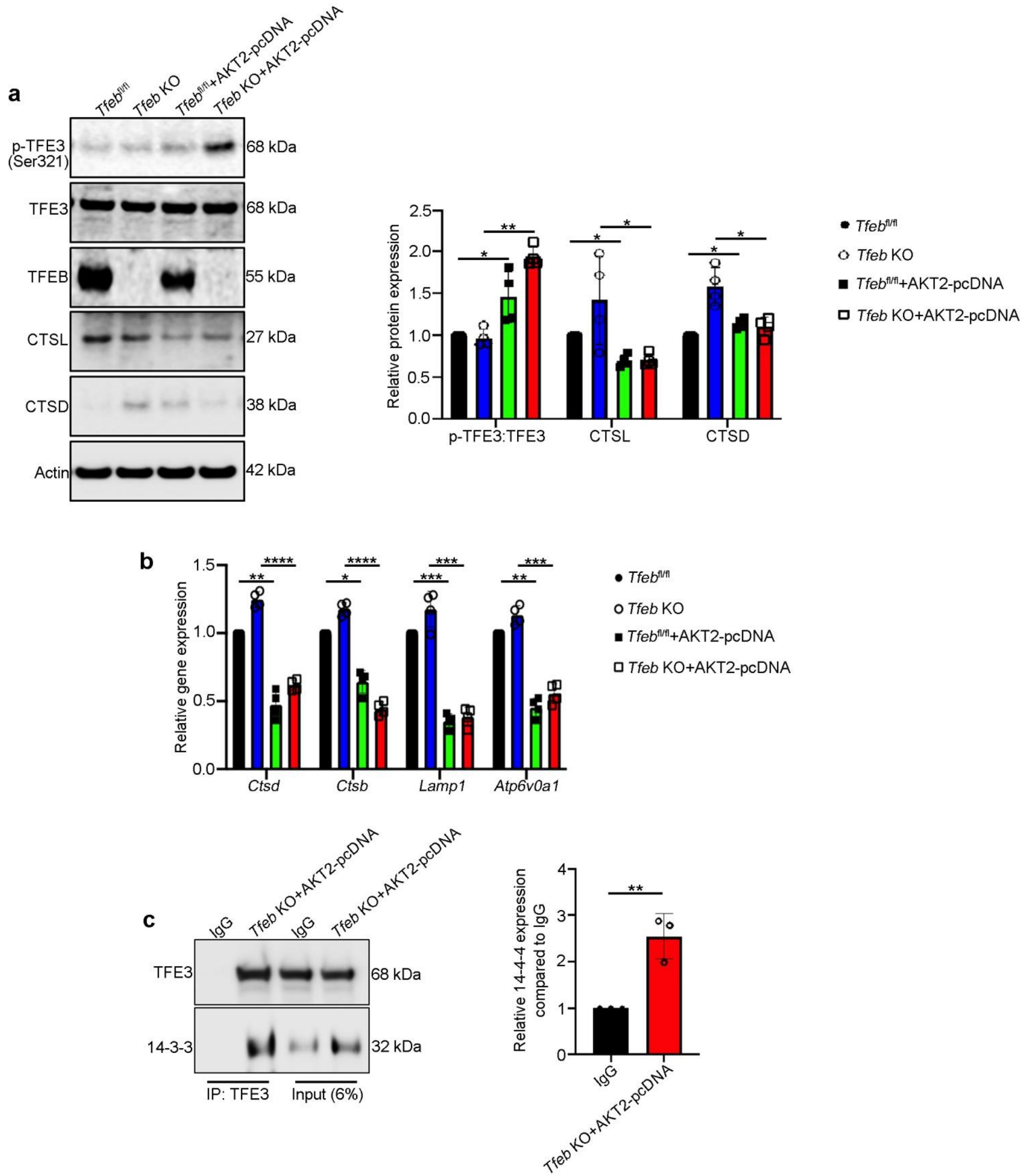


**<u>Supplementary Figure 1</u>: *Akt2 inhibits TFE3-mediated compensatory lysosomal function upon loss of TFEB*.** (**a**) Western blot analysis showing that Akt2 overexpression in starved (20 h in serum-free medium) *Tfeb* KO and *Tfeb* floxed mouse embryonic flibroblasts (MEFs), increased the protein levels of p-TFE3 (S321) and decreased CTSD and CTSL, compared to control (untransfected cells). n=4. (**b**) Expression of *Ctsd*, *Ctsb*, *Lamp1*, and *Atp6v0a1* also showed noticeable downregulation in Akt2 overexpressing and starved *Tfeb* KO and *Tfeb* floxed MEFs compared to controls. n=4. (**c**) Co-immunoprecipitation studies (with pulling down of TFE3 and immunoblotting for 14-3-3) showing that in Akt2 overexpressing *Tfeb* KO MEFs, TFE3 binds more to 14-3-3, as compared to IgG pull-down controls, perhaps accounting for its proteasomal degradation and subsequent inhibition of nuclear translocation. n=4. ****P<0.0001, ***P<0.001, **P<0.01, *P<0.05.

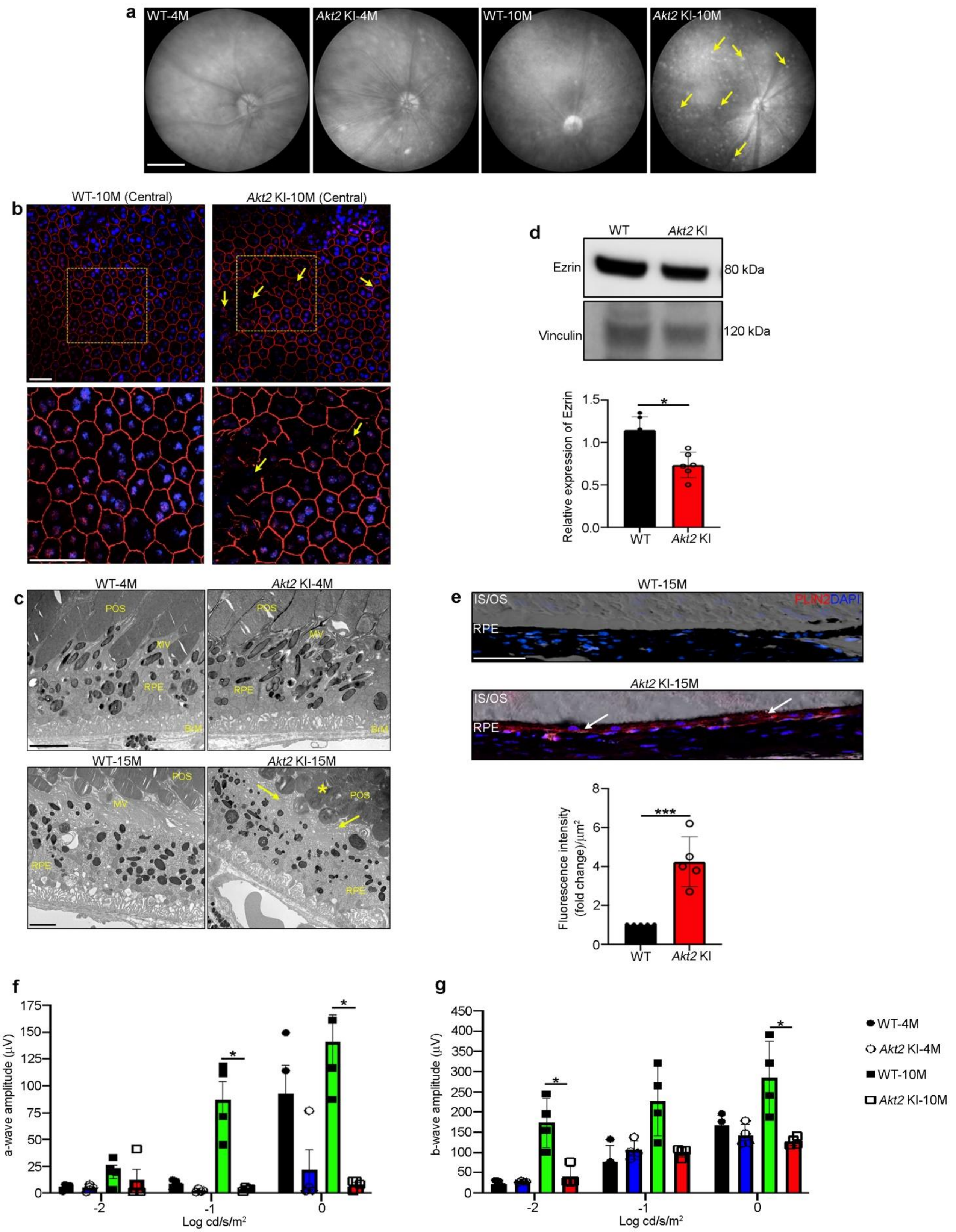


**Supplementary Figure 2**: ***Akt2 KI mice develop an AMD-like phenotype***. (**a**) Fundus photographs showing accumulation of autofluorescent foci (arrows in **a**) in 10-month-old *Akt2*

KI mice, but not in age-matched controls or young animals. n=5. (**b**) Immunostaining with ZO-1 on RPE flatmount from 10-month-old *Akt2* KI mice showing alterations in the normal cobblestone-like morphology in the central retinal region (arrows in **b**), not seen in age-matched WT. n=4. Scale bar= 50 μm (Zoomed Inset= 80 μm). (**c**) Transmission electron micrographs showing loss of microvilli (MV; arrows in *Akt2* KI) and abnormal photoreceptor outer segments (POS; asterisk in *Akt2* KI) in 15-month-old *Akt2* KI RPE cells, but not in age-matched WT or young (4 month old) *Akt2* KI mice. n=5. Scale bar= 2 μm. (**d**) Western blot showing decreased expression of ezrin in 10-month-old *Akt2* KI RPE cells, relative to control (WT). n=4. (**e**) Immunofluorescence studies showing increased accumulation of PLIN2 (red; arrows in *Akt2* KI) in 15-month-old *Akt2* KI RPE, compared to WT. n=5. Scale bar= 50 μm. (**f**,**g**) Electroretinography analysis showing decrease in scotopic (**f**) a-wave and (**g**) b-wave amplitudes in 10- month- old *Akt2* KI mice, compared to controls. Such changes were not seen in young (4 month old) mice. n=4. All values are Mean ± S.D. ***P<0.001, *P<0.05.

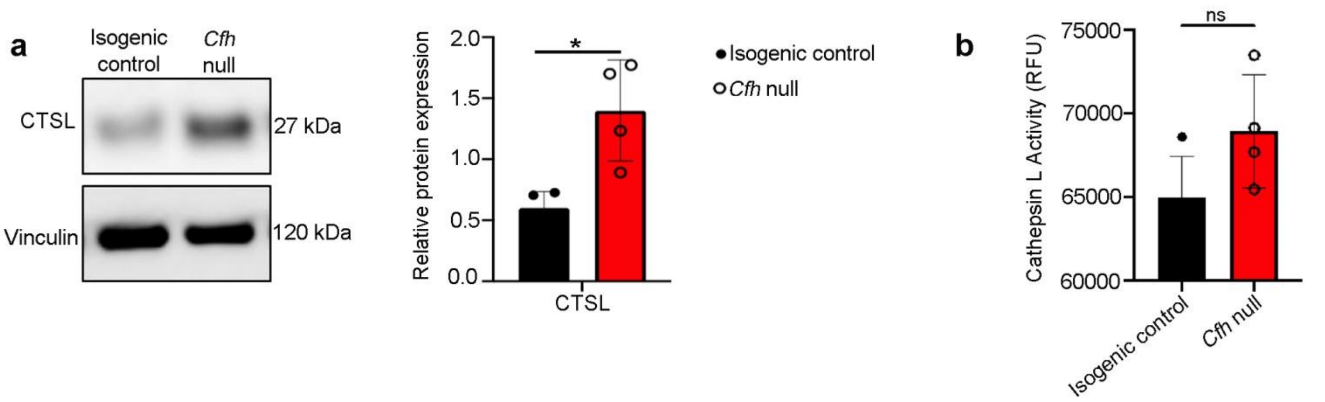


<u>Supplementary Figure 3</u>: ***Loss of CFH does not result in abnormal lysosomal function in CFH null cells***. (**a**) Western blot and densitometric analysis showing no significant change in Cathepsin L (CTSL) protein levels or (**b**) activity in *CFH* null (*CFH*$^{-/-}$) iRPE cells, compared to isogenic controls. n=4. *P<0.05. ns= non-significant.

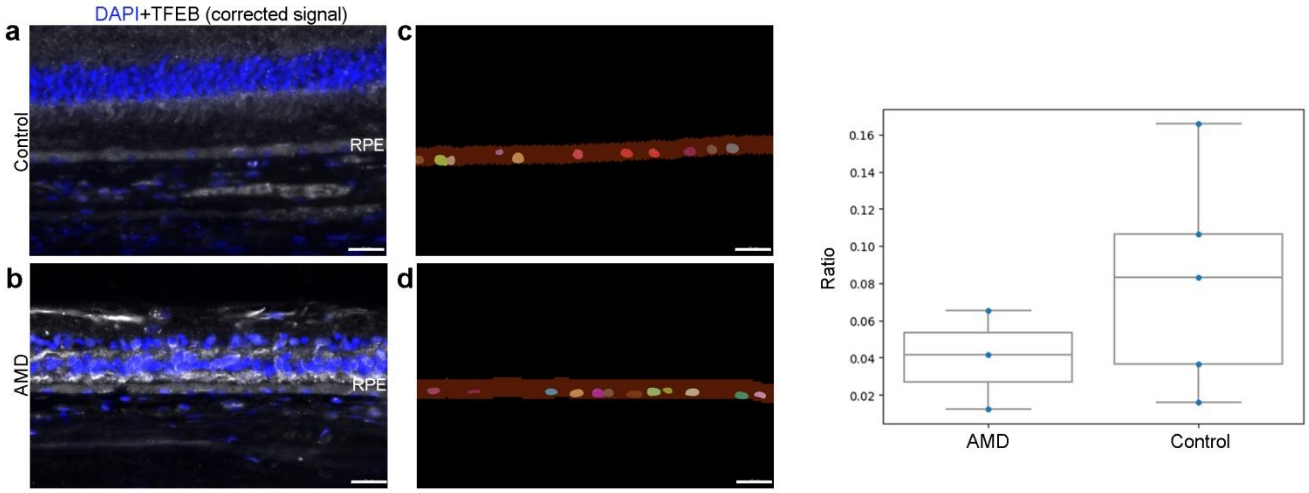


**<u>Supplementary Figure 4</u>**: ***TFEB expression in human AMD***. Immunohistochemistry followed by AI-based quantification on human donor retinal sections showing a trend towards decrease in the presence of TFEB nuclear foci (white) in (**b**) AMD patients, compared to (**a**) controls. (**c**, **d**) Detection of RPE layer (brown), individual nuclei (multicolors) and TFEB foci (white). Nuclei were considered positive when containing at least two TFEB foci. Ratio of positive over all detected nuclei is presented in the boxplot. Scale bar= 25 μm. Control (n=5) and AMD (n=3).

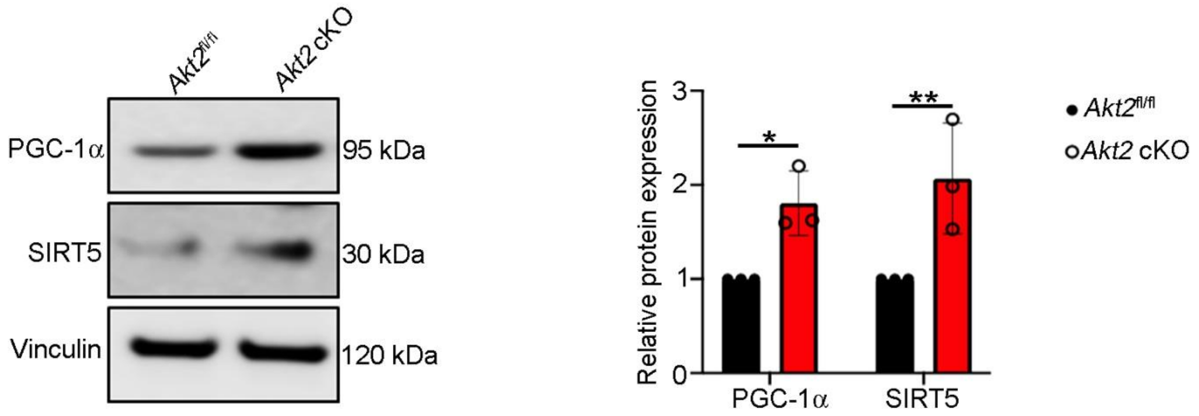


**Supplementary Figure 5**: ***Akt2 cKO RPE cells show upregulation in PGC-1α and SIRT5 levels***. Western blot showing elevated levels of both PGC-1α and SIRT5 in *Akt2* cKO RPE cells, relative to floxed controls (*Akt2*$^{fl/fl}$). n=3. All values are Mean ± S.D. **P<0.01, *P<0.05.

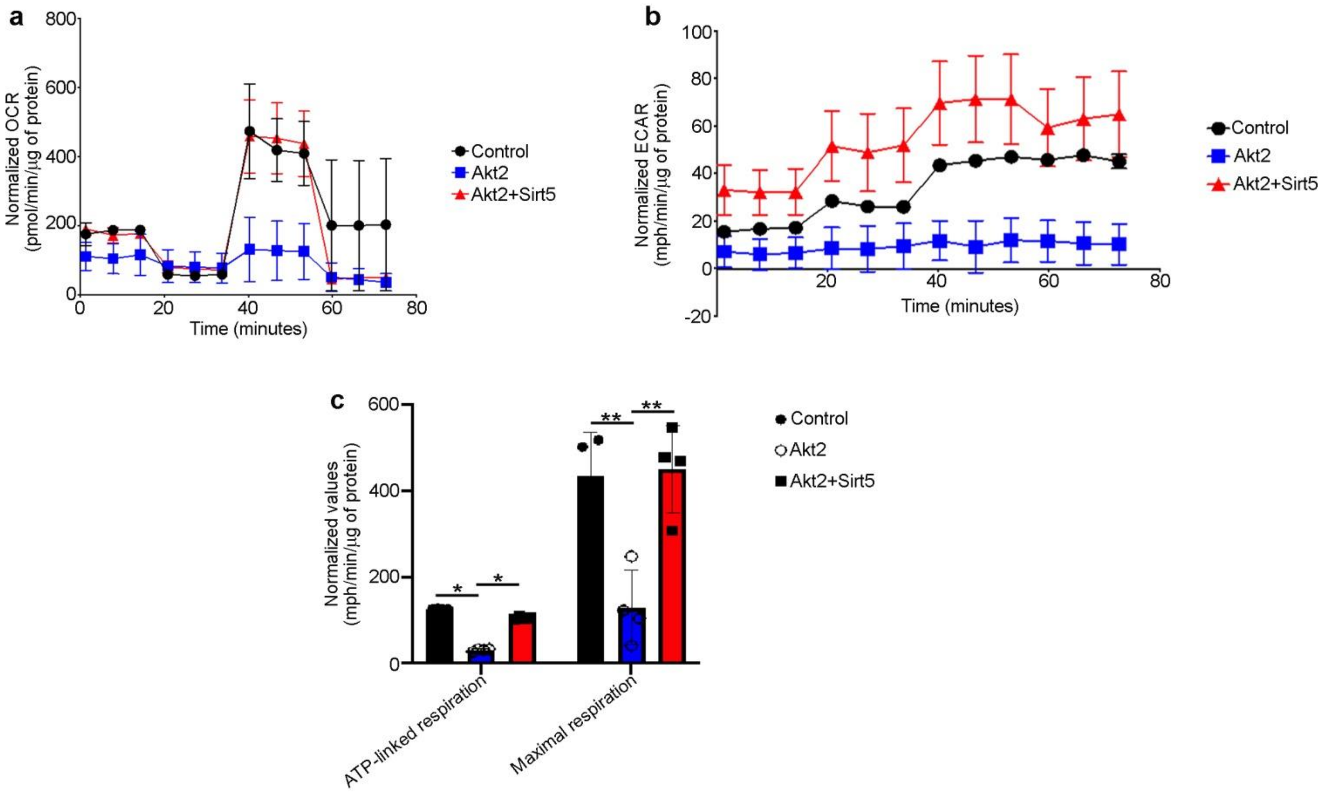


**<u>Supplementary Figure 6</u>**: ***Alterations in mitochondrial function in RPE cells overexpressing AKT2***. Seahorse analysis using the mitostress assay showing time-dependent changes in metabolic flux. Plots show (**a**) oxygen consumption rate (OCR) and (**b**) extracellular acidification rate (ECAR) upon treatment with mitochondrial respiration blockers oligomycin, Carbonyl cyanide-p-trifluoromethoxy phenylhydrazone (FCCP), and Rotenone/Antimycin A at particular time points in untransfected ARPE19 cells (control) or cells overexpressing AKT2 or cells overexpressing both AKT2 and SIRT5. n=4. (**c**) ATP-linked respiration and maximal respiration were significantly decreased in ARPE19 cells overexpressing AKT2, compared to controls. Such changes were rescued in cells overexpressing both AKT2 and SIRT5, indicating that SIRT5 can inhibit AKT2-mediated mitochondrial alterations. n=4. All values are Mean ± S.D. **P<0.01, *P<0.05.

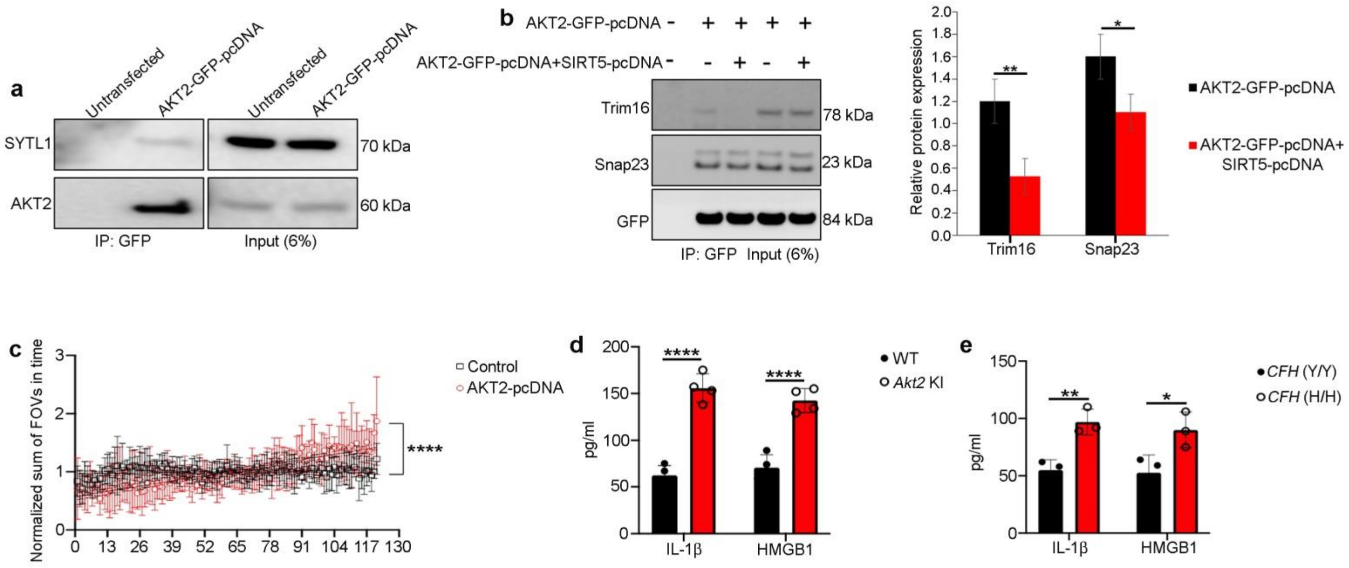


**<u>Supplementary Figure 7</u>**: ***Akt2 upregulation is associated with activation of secretory autophagy in the RPE cells.*** (**a**) Co-immunoprecipitation studies showing binding of SYTL1 to AKT2 in ARPE19 cells overexpressing AKT2-GFP when AKT2 was pulled down using anti-GFP magnetic beads. n=3. (**b**) Pulldown assay with anti-GFP magnetic beads from lysates of ARPE19 cells overexpressing either GFP-AKT2 or GFP-AKT2 and SIRT5-HA and serum/nutrient starved for 1h in HBSS. Cells overexpressing only GFP-AKT2 showed an increase in binding of secretory autophagy mediators Snap23 and Trim16. This binding was reduced upon simultaneous overexpression of SIRT5. n=4. (**c**) Linear regression plot showing significant change in autophagosome number on the cell membrane as evident from TIRF microscopy in AKT2 overexpressing (Akt2-pcDNA transfected) ARPE19 cells, compared to untransfected controls. n=3. ELISA showing increased levels of IL-1β and HMGB1 in spent medium from cultured (**d**) *Akt2* KI RPE explants and (**e**) iPSC-derived RPE cells from *CFH* Y402H risk allele containing donors [*CFH* (H/H)], compared to controls. n=3. All values are Mean ± S.D. ****P<0.0001, **P<0.01, *P<0.05.

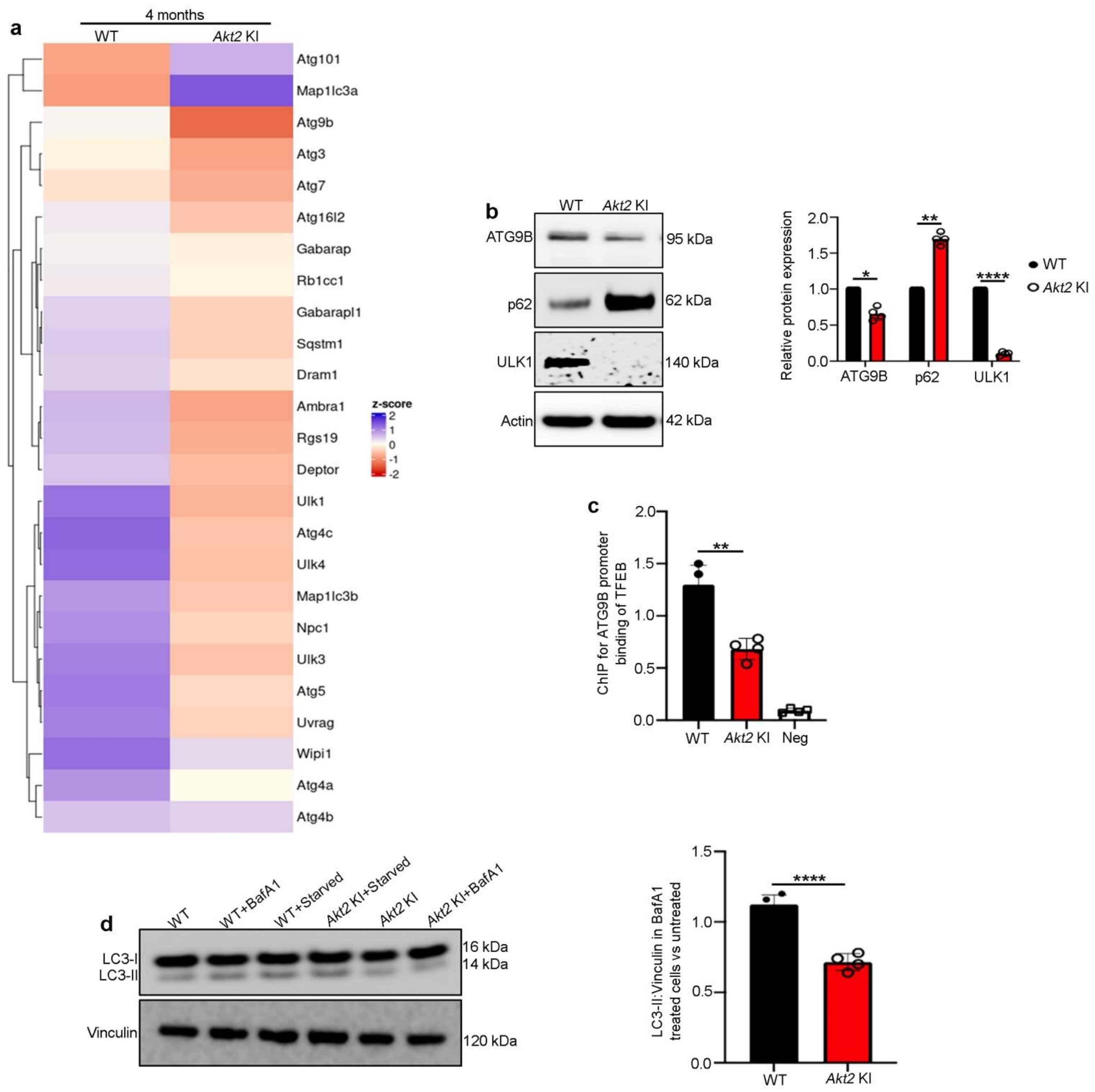


**<u>Supplementary Figure 8</u>**: ***Akt2 KI RPE cells show diminished autophagy***. (**a**) RNAseq analysis from RPE cells of 4-month-old WT and *Akt2* KI mice showing differential expression of several autophagy related genes. n=3. (**b**) Western blot showing reduced expression of autophagy mediators ATG9B and ULK1, as well as upregulation of the autophagosome marker p62/SQSTM1 in *Akt2* KI RPE cells, relative to WT. n=4. (**c**) Chromatin immunoprecipitation showing diminished binding of TFEB on Atg9b promoter region in *Akt2* KI RPE cells, compared to WT. n=4. (**d**) Western blot showing reduced autophagy flux (Ratio of LC3-II/ Vinculin in BafA1 treated vs untreated) in *Akt2* KI RPE explants compared to WT when treated with Bafilomycin A1 (BafA1; 1 $\mu$m) for 4 h. n=4. All values are Mean ± S.D. ****P<0.0001, **P<0.01, *P<0.05.

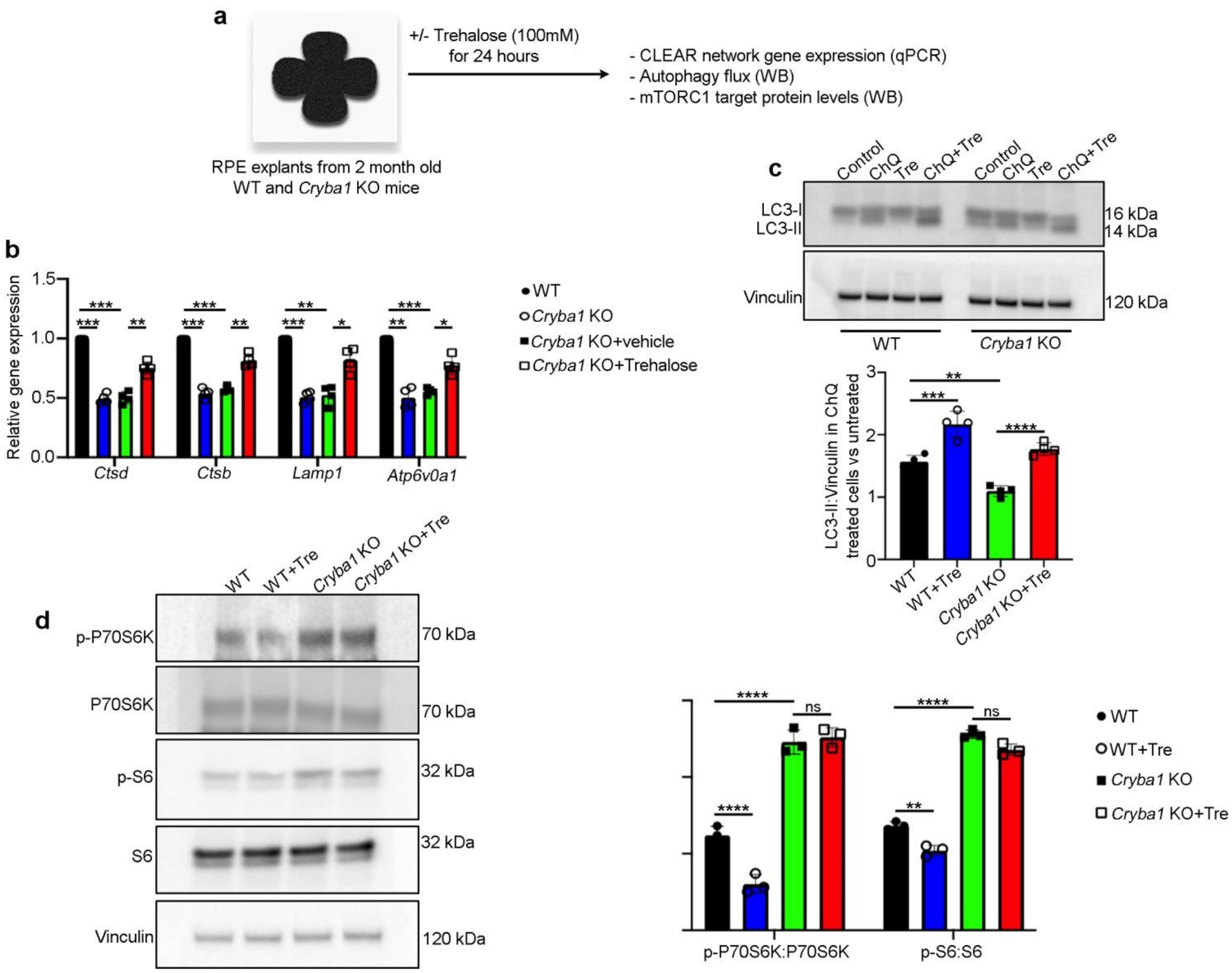


<u>**Supplementary Figure 9**</u>: ***Trehalose rescues lysosomal and autophagy abnormalities in Cryba1 KO RPE explants***. (**a**) Schematic showing experimental design to evaluate efficacy of trehalose in rescuing autophagy and lysosomal abnormalities in *Cryba1* KO RPE explants. (**b**) qPCR showing trehalose (100 mM for 24 h) treatment of *Cryba1* KO RPE explants could rescue the expression levels of CLEAR network genes like *Ctsd*, *Ctsb*, *Lamp1* and *Atp6v0a1*. n=4. (**c**) Western blot showing trehalose treatment could rescue the decrease in autophagy flux (Ratio of LC3-II/ Vinculin in ChQ treated vs untreated) in *Cryba1* KO RPE explants when treated with chloroquine (ChQ; 50 $\mu$m) for 6 h, compared to untreated *Cryba1* KO RPE explants. n=3. (**d**) Western blot showing no noticeable difference in mTORC1 downstream

mediators P70S6K and S6 in *Cryba1* KO RPE explants treated with trehalose, indicating that the effect of trehalose is independent of mTORC1 signaling. n=3. All values are Mean ± S.D. ****P<0.0001, ***P<0.001, **P<0.01, *P<0.05. ns=not significant.

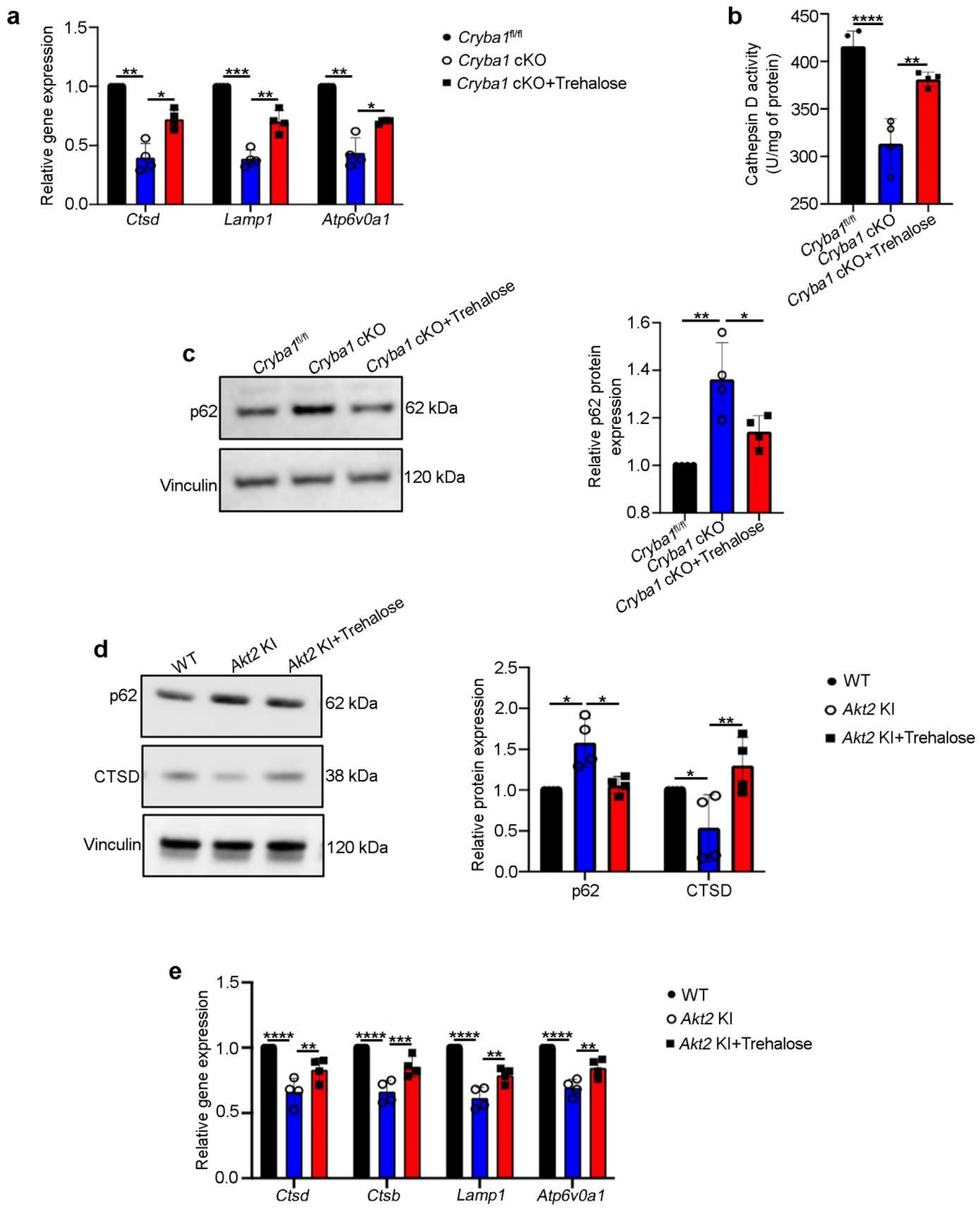


**<u>Supplementary Figure 10</u>**: ***Trehalose rejuvenates lysosomal function and rescues autophagy in mouse models in vivo***. RPE cells from 6-month-old *Cryba1* cKO mice treated with trehalose for 3 consecutive months show significant rescue in the levels of (**a**) CLEAR network genes like *Ctsd*, *Lamp1* and *Atp6v0a1*, (**b**) cathepsin D (CTSD) activity and (**c**) protein levels of the autophagosome marker p62/SQSTM1, relative to RPE cells from vehicle (water) treated *Cryba1* cKO mice. n=4. RPE cells from 6-month-old *Akt2* KI mice treated with trehalose for 3 consecutive months showed statistically significant rescue of (**d**) protein levels of autophagy mediator ATG9B and autophagosome marker p62/SQSTM1, as well as CLEAR network gene (*Ctsd*, *Ctsb*, *Lamp1*, *Atp6v0a1*) expression, compared to vehicle treated *Akt2* KI mice. n=4. All values are Mean ± S.D. ****P<0.0001, ***P<0.001, **P<0.01, *P<0.05.

| Name | ID | Z Score | S Score | F635 | B635 |
|---|---|---|---|---|---|
| SIRT5 | JHU04487.B1C25R68 | 66.32 | 34.394 | 1576.5 | 47 |
| SORBS3 | JHU04395.B1C9R68 | 31.926 | 9.929 | 788.5 | 48.5 |
| MAB21L1 | JHU04362.B2C17R68 | 21.997 | 1.244 | 561 | 48 |
| HIST1H1A | JHU10506.B8C29R78 | 20.753 | 4.147 | 532.5 | 48.5 |
| H1FX | JHU08674.B6C11R48 | 16.606 | 0.087 | 437.5 | 49.5 |
| PPP1R3B | JHU05921.B5C14R2 | 16.519 | 0.022 | 435.5 | 46.5 |
| SCL-70 | Auto-antigen.B20C6R42 | 16.497 | 0.109 | 435 | 49 |
| CXorf51B | JHU10867.B5C14R84 | 16.388 | 0.589 | 432.5 | 48 |
| CYB561 | JHU13134.B9C27R28 | 15.799 | 0.000 | 419 | 48 |
| SYTL1 | JHU08341.B8C4R38 | 15.799 | 1.332 | 419 | 47.5 |
| DIMT1 | JHU09043.B8C6R54 | 14.467 | 0.392 | 388.5 | 47.5 |
| PNKP | JHU08225.B5C9R42 | 14.075 | 0.284 | 379.5 | 47 |
| C7orf50 | JHU08406.B8C21R38 | 13.791 | 0.066 | 373 | 48 |
| THYN1 | JHU16333.B10C4R76 | 13.725 | 0.000 | 371.5 | 48.5 |
| JHU04032 | JHU04032.B3C6R62 | 13.725 | 0.589 | 371.5 | 48.5 |
| Lupus La | Auto-antigen.B20C4R38 | 13.136 | 0.458 | 358 | 49 |
| SRSF5 | JHU04389.B9C3R88 | 12.678 | 0.044 | 347.5 | 54.5 |
| VRK1 | JHU10925.B7C27R80 | 12.634 | 0.218 | 346.5 | 48 |
| ANXA2 | JHU13600.B12C25R36 | 12.416 | 0.502 | 341.5 | 48.5 |
| KHDRBS1 | JHU15238.B9C18R60 | 11.914 | 0.065 | 330 | 47 |
| GYS1 | JHU08863.B5C3R52 | 11.849 | 0.022 | 328.5 | 48 |

**Supplementary Table 1**: ***AKT2 binding partners***. Human high-throughput protein-protein interaction study showing several AKT2 binding partners with their Z-scores. SIRT5 and SYTL1 are highlighted with red and blue, respectively.

**Supplementary Movies 1 and 2**: ***TIRF imaging for autophagosome binding to cell surface***. Time lapse movies from control (Supplementary Movie 1) and Akt2 overexpressing (Supplementary Movie 2) ARPE19 cells showing GFP-LC3 puncta at the cell surface over as a function of time, upon induction of starvation (incubation in serum free medium). n=3.
